# Size control of the inner ear via hydraulic feedback

**DOI:** 10.1101/349381

**Authors:** Kishore R. Mosaliganti, Ian A. Swinburne, Chon U Chan, Nikolaus D. Obholzer, Amelia A. Green, Shreyas Tanksale, L. Mahadevan, Sean G. Megason

## Abstract

Animals make organs of precise size, shape, and symmetry despite noise in underlying molecular and cellular processes. How developing organs manage this noise is largely unknown. Here, we combine quantitative imaging, physical theory, and physiological measurement of hydrostatic pressure and fluid transport in zebrafish to study size control of the developing inner ear. We find that fluid accumulation creates hydrostatic pressure in the lumen leading to stress in the epithelium and expansion of the otic vesicle. Pressure, in turn, inhibits fluid transport into the lumen. This negative feedback loop between pressure and transport allows the otic vesicle to change growth rate to control natural or experimentally-induced size variation. Spatiotemporal patterning of contractility modulates pressure-driven strain for regional tissue thinning. Our work connects molecular-driven mechanisms, such as osmotic pressure driven strain and actomyosin tension, to the regulation of tissue morphogenesis via hydraulic feedback to ensure robust control of organ size.

## HIGHLIGHTS

- Hydrostatic pressure in the lumen drives viscoelastic growth of the otic vesicle
- Mathematical model links growth with geometry, fluid flux, and tissue viscoelasticity
- A negative feedback loop between pressure and fluid flux controls inner ear size
- Spatiotemporal patterning of actomyosin contractility results in regional differences in tissue stretching

## ETOC

Most internal organs—including the eyes, lung, gut, kidney, bladder, brain, and vascular system—all begin as epithelialized cysts or tubes with a fluid-filled lumen. Using the inner ear, we show that fluid transports creates hydrostatic pressure in the lumen that drives growth, while negative feedback between pressure and fluid flux ensures control of organ size. Furthermore, the shape of the inner ear is modulated by spatiotemporal patterning of actomyosin contractility allowing a uniform lumenal pressure to drive varying epithelial strain rates. Mathematical modeling illuminates how organ size and shape control occurs by integrating molecular signals with growth, geometry and mechanics.

## INTRODUCTION

A fundamental question in developmental biology is how different organs acquire their proper sizes, which are necessary for their healthy function. The existence of control mechanisms is evident in the consistency of organ size in the face of intrinsic noise in biological reactions such as gene expression, and in the observed recovery from size perturbations during development (Waddington, 1959; Debat and Peronnet, 2013; Rao et al., 2002; Lestas et al., 2010). However, unlike in engineered systems, where there is often a clear distinction and hierarchy between the controller and the system, in organ growth one may not have a clear hierarchy—instead there may be control mechanisms distributed across tissues and across scales. Furthermore, in developmental biology, we observe an evolved system that is not necessarily robust to all experimental perturbations that we apply when trying to understand their control networks. Consequently, it can be difficult to distinguish what is necessary for growth from what controls size.

Identifying specific mechanisms that coordinate growth—to ultimately control organ size—has been difficult because the phenomenon of growth encompasses regulatory networks that can span the molecular to organismic. Classical organ transplantation and regeneration studies in the fly (Bryant and Levinson, 1985; Hariharan, 2015), mouse (Metcalf, 1963; Metcalf, 1964), and salamander (Twitty and Schwind, 1931) have indicated that both organ-autonomous and non-autonomous mechanisms control size. In his “chalone” model, Bullough’s proposed growth duration to be regulated by an inhibitor of proliferation that is secreted by the growing organ and upon crossing a concentration threshold stops organ growth at the target size (Bullough and Laurence, 1964). Modern evidence for organ intrinsic chalones exists in myostatin for skeletal muscle, GDF11 for the nervous system, BMP3 for bone, and BMP2/4 for hair (McPherron et al., 1997; Wu et al., 2003; Plikus et al., 2008; Gamer et al., 2009). Several existing models for size control are based on global positional information regulating cell proliferation based on a morphogen gradient until final organ size is achieved (Day and Lawrence, 2000; Rogulja and Irvine, 2005; Wartlick et al., 2011). Other models emphasize the role of local cell-cell interactions in regulating cell proliferation or cell lineages to make tissues of the correct proportions (García-Bellido, 2009; Kunche et al., 2016). Given that cells are coupled to each other through cell-cell and cell-substrate contacts, physical constraints and tissue geometry provide tissue-level feedback. More recent models emphasize the role of tissue mechanics in regulating cell proliferation via anisotropic stresses and strain rates (Shraiman, 2005; Ingber, 2005; Savin et al., 2011; Hufnagel et al., 2007; Behrndt et al., 2012; Irvine and Shraiman, 2017; Nelson et al., 2017; Pan et al., 2016). From a molecular perspective, the insulin, Hippo (Dupont et al., 2011; Legoff et al., 2013; Pan et al., 2016) and TOR signaling pathways (Colombani et al., 2003; Zhang et al., 2000) have been well-established as regulators of organ size. Several studies have demonstrated that genetic mutation in these pathways is sufficient to alter organ or body size through increases in cell number, cell size, or both (Tumaneng et al., 2012), but the mechanisms that control size in the engineering sense (e.g. feedback of size on growth rate) are generally not known.

Most size control theories have focused on regulation of cell proliferation. Control may also arise from regulation of other parameters such as cell shape, material properties, transepithelial transport, adhesion, and the extracellular environment. In particular, fluid accumulation is a feature of developmental growth for several luminized organs including the embryonic brain (Desmond and Jacobson, 1977; Lowery and Sive, 2005), eye (Coulombre, 1956), gut (Bagnat et al., 2007), Kupffer’s vesicle (Navis et al., 2013; Dasgupta et al., 2018), and inner ear (Abbas and Whitfield, 2009; Hoijman et al., 2015). Water transport across an epithelium underlies these phenomenon (Frömter and Diamond, 1972; Günzel and Yu, 2013; Rubashkin et al., 2005; Fischbarg, 2010), and for the developing brain and eye it was shown that fluid accumulation coincides with increased hydrostatic pressure (Desmond and Jacobson, 1977; Coulombre, 1956). Although specific ion transporters necessary for fluid accumulation have been identified (Lowery and Sive, 2005; Bagnat et al., 2007; Navis et al., 2013; Abbas and Whitfield, 2009), it is currently unclear how ion transport and transepithelial fluid flow are regulated, or what their role in growth control is.

Catch-up growth during development is the phenomenon where, after growth delay or perturbation, an organ transiently elevates its growth rate relative to other organs to get back on course. During fly development, if the growth of one imaginal disc is perturbed then a hormone, ecdysone, signals to the other imaginal discs to slow their growth such that the perturbed organ can catch-up and the animal’s coordinated growth can resume (Parker and Shingleton, 2011). The phenomenon of catch-up growth clarified ecdysone’s activity as being important for size control. Catch-up growth also occurs in vertebrates: if an infant heart or kidney is transplanted into an adult, it grows faster than the surrounding tissue to catch-up to a target size (Dittmer et al., 1974; Silber, 1976). Catch-up growth also occurs during bone development (Rosello-Diez and Joyner, 2015). Recently, the related phenomenon of organ symmetry has been addressed in the context of long bones, tails, and the inner ear; but, the control mechanism underlying catch-growth was not clearly identified (Rosello-Diez et al., 2017; Das et al., 2017; Green et al., 2017). While catch-up growth has been successfully used in the study of insects to discover an organ size control mechanism, catch-up growth had yet to be leveraged to discover a vertebrate specific mechanism of organ size control.

Here we use a newly revealed instance of catch-up growth combined with physical theory to uncover how size control is achieved in the zebrafish otic vesicle, a fluid-filled closed epithelium that develops into the inner ear. We postulate that fluid pressure is a fundamental regulator of developmental growth in lumenized organs and hydraulic feedback can give rise to robust control of size.

## RESULTS

### *In toto* imaging of otic vesicle development shows lumenal inflation dominates growth, not cell proliferation

We sought to determine how size control is achieved in the zebrafish otic vesicle, a 3D lumenized epithelial cyst that becomes the inner ear. Prior studies used qualitative observations and 4D imaging to examine the formation of the otic vesicle (Haddon and Lewis, 1996; Hoijman et al., 2015; Dyballa et al., 2017). To systematically investigate inner ear morphogenesis at longer timescales between 12-45 hpf, we used high-resolution 3D+*t* confocal imaging combined with automated algorithms for quantifying cell and tissue morphology (Figure S1A-F) (Megason, 2009). Beginning at 12 hours post-fertilization (hpf), bilateral regions of ectoderm adjacent to the hindbrain proliferate and subcutaneously accumulate to form the otic placodes (Movie S1, Figure 1A). The complex morphology of the inner ear arises from progressive changes in cell number, size, shape, and arrangement along with tissue-level patterns of polarization (12-14 hpf, Figure 1A), mesenchymal-to-epithelial transition (14-16 hpf, Figure 1B) and cavitation (16-24 hpf, Figure 1C). These steps build a closed ovoid epithelial structure, the otic vesicle, filled with a fluid called endolymph (Figure S1 G-H). After assembly, the otic vesicle undergoes a period of rapid growth (16-45 hpf, Figure 1D) prior to the development of more complex substructures such as the semicircular canals and endolymphatic sac.

**Figure 1:**
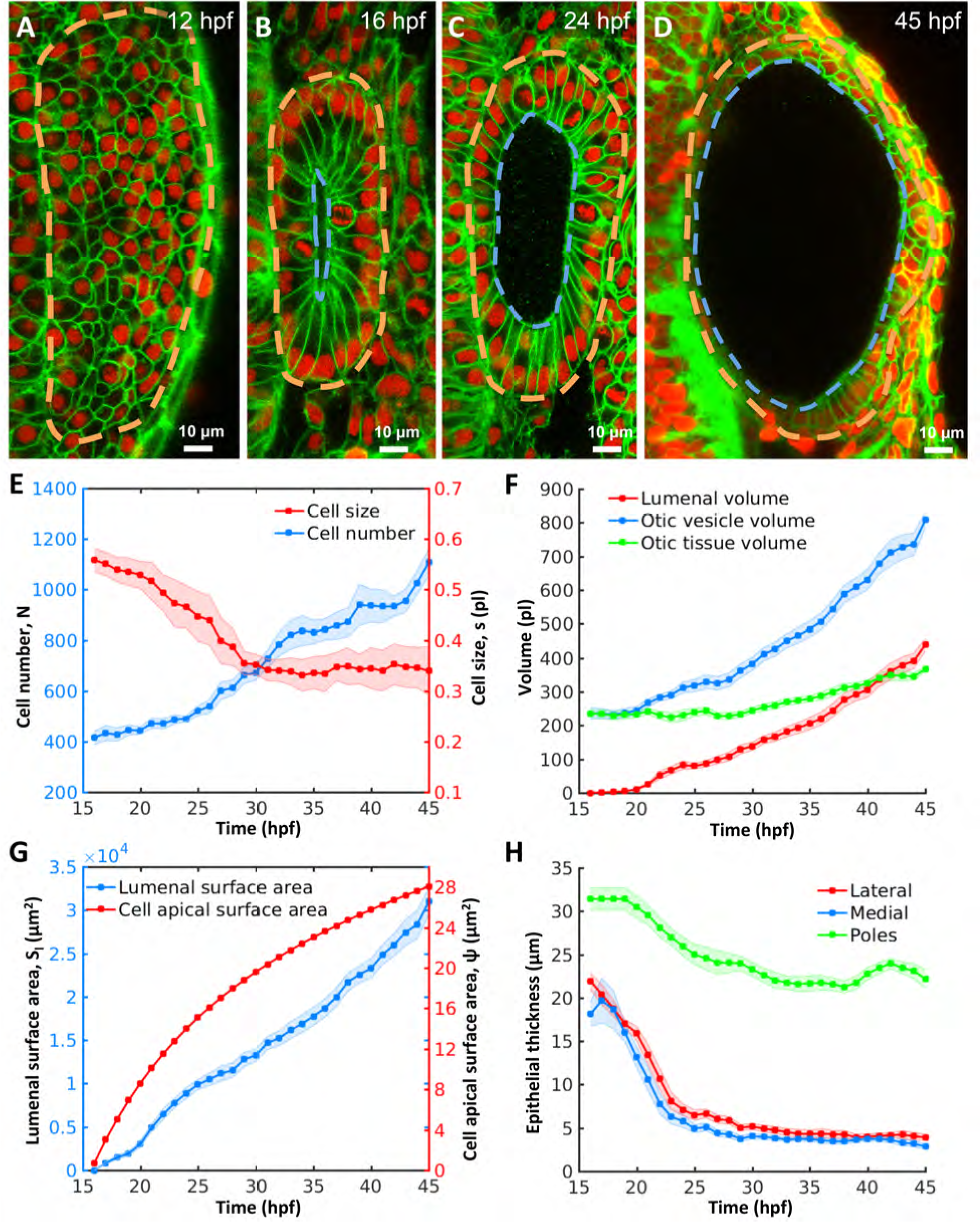
Morphodynamic analysis of inner ear growth from 16-45 hpf using *in toto* imaging. **(A-D)** Confocal micrographs of otic vesicle development at (A) 12 hpf, (B) 16 hpf, (C) 24 hpf, and (D) 45 hpf. Orange and blue contours demarcate otic vesicle and lumenal surfaces respectively. Embryos are double transgenic for highlighting membranes and nuclei (*Tg(actb2:Hsa.H2B-tdTomato)*^*hm25*^; *Tg(actb2:mem-citrine)*^*hm26*^). n = 10 embryos per datapoint. Error bars are SD. **(E)** Primary y-axis plots cell numbers (*N*, blue markers) and secondary axis plots average cell size (*s*, picoliters or pl, red markers). **(F)** Quantification of vesicle (*V*_*o*_, blue markers), lumenal (*V*_*l*_, red markers), and tissue volumes (*V*_*t*_, green markers). **(G)** Primary y-axis plots lumenal surface area (*S*_*l*_ blue markers). Secondary axis plots average cell apical surface area (*s*, red markers) evaluated numerically by fitting quadratic polynomials to surface area (*S*_*l*_) and cell number (*N*) data. **(H)** Quantification of wall thickness (*h*, μm) at locations next to the hindbrain (medial, blue), ectoderm (lateral, red), and anterioposterior poles (poles, green). Related to Figure S1 and Movie S1.

To evaluate growth kinetics, we used 3D image analysis (Figure S1 I-M) to quantify a number of morphodynamic parameters between 16 and 45 hpf. During this period, cell number increased nearly three-fold from 415±26 to 1106±52 cells (blue curve, Figure 1E). However, cell proliferation was offset by a decrease in average cell size from 0.55±0.02 pl at 16 hpf to 0.34±0.03 pl at 28 hpf and stayed constant thereafter (red curve). Tissue volume, the product of cell number and average cell size, remained effectively constant (230.6±7.4 pl) until 28 hpf and subsequently increased linearly by 132 pl to 45 hpf (green curve, Figure 1F). The volume of the otic vesicle increased dramatically, by 572 pl from 235±16 pl to 807±23 pl (blue curve). The majority of the increase in size of the otic vesicle is due to an increase in lumen volume (red curve) from 0 to 440±18 pl (77% of the total increase) while tissue growth contributed only 23% to the increase in size.

### Pressure inflates the otic vesicle and stretches tissue viscoelastically

A mismatch between the volumetric growth of the lumen and the tissue enclosing the lumen indicates a potential role for otic tissue remodeling. Since the cube root of lumen volume increases more rapidly than the square root of the cell number enclosing that volume, we investigated how epithelial cell shape changes to accommodate growth. We observed a monotonic increase in average cell apical surface area (*ψ* = *S*_*l*_/*N*; *S*_*l*_ is lumenal surface area, *N* is the cell number, Figure 1G). Since the otic epithelium is not uniform in thickness, we examined regions of the epithelium that contribute to the stretch. Except for the future sensory patches at the anterior and posterior ends (poles), epithelial thickness of the remaining otic vesicle significantly decreased from 20 μm to 4 μm during otic vesicle growth (lateral and medial regions, Figure 1H). The increase in lumenal volume and large cell stretching rates suggested that the vesicle is pressurized and the epithelium is under tension.

In the absence of extrinsic forces, cells round-up to a spherical morphology during mitosis by balancing internal osmotic pressure with tension provided by cortical actomyosin (Stewart et al., 2011). To investigate the development of pressure derived stress in the epithelium, we used mitotic cells as “strain gauges” by measuring their deviation in aspect ratios from spheres. We observed that mitotic cells fail to round up fully in regions where the otic epithelium pushes against the hindbrain and ectoderm, in contrast to non-contact regions at the anterioposterior poles (Figure 2A,C-G). Furthermore, cell division planes are closely aligned with the surface-normal to the epithelium (blue markers, Figure 2H) in comparison to the broader distribution exhibited by the non-contact cell populations (red). The overall alignment progressively increases in developmental time as cells become more stretched (Figure 2I), consistent with mechanical stress driven spindle alignment previously observed in various systems including the zebrafish gastrula (Campinho et al., 2013), fly imaginal disc (Legoff et al., 2013), and zebrafish pre-enveloping layer (Xiong et al., 2014). Given that the otic vesicle is wedged between the hindbrain and skin (Figure S1G-H), we examined the impact of its volumetric growth on these tissues. We reasoned that if pressure is present, the vesicle would exhibit higher rigidity and consequently deform neighboring structures as it increased in size. To test this idea, we quantified the indentation of the hindbrain and otic vesicle interface. We observed that as the vesicle grows, the initially planar hindbrain surface indents in, and the skin bulges out (Figure 2B,C-F,J).

**Figure 2:**
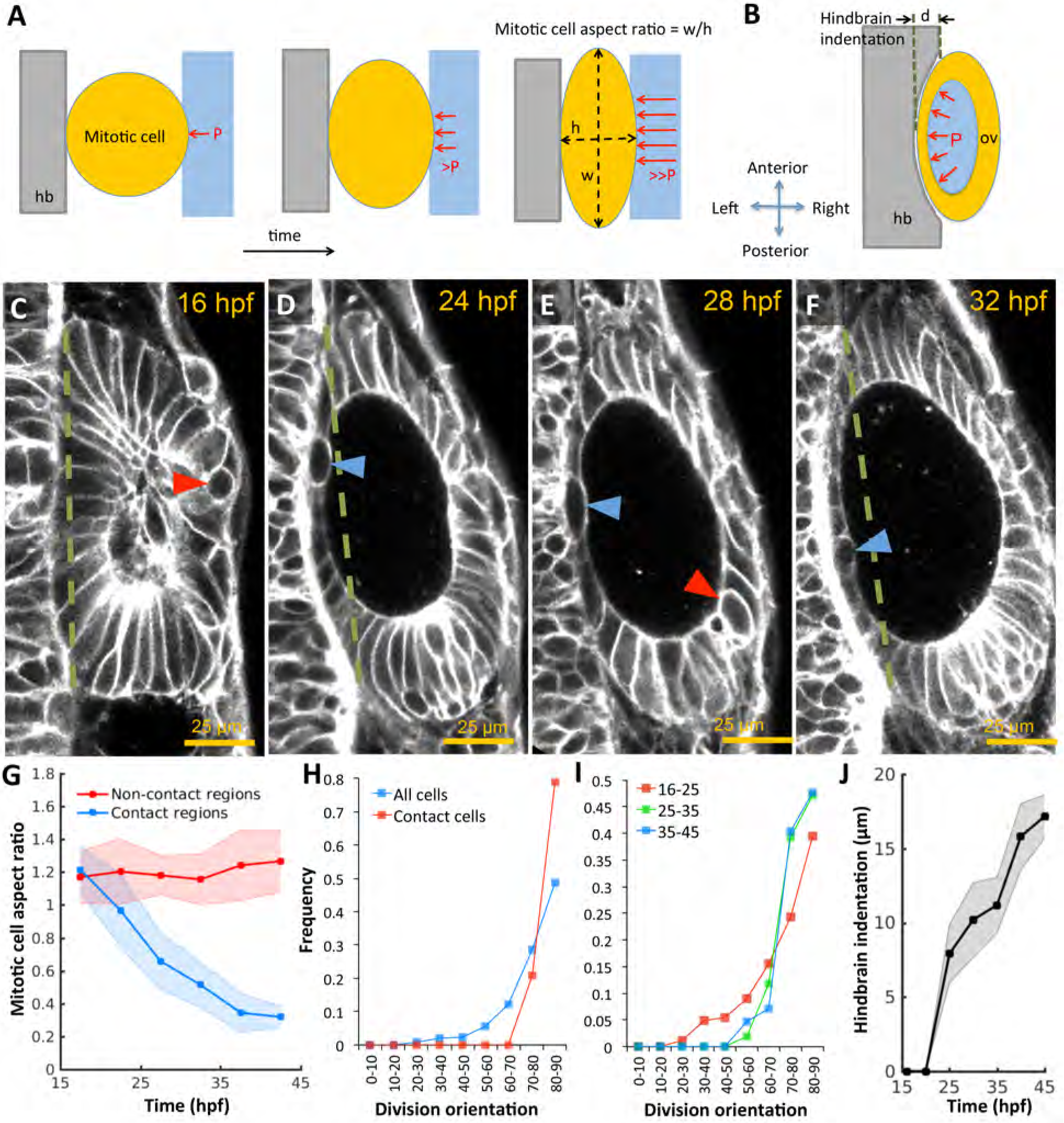
Otic vesicle growth is correlated with deformations in mitotic cell shapes and neighboring tissues that are indicative of pressure-driven strain. **(A)** Diagram illustrating inhibition of mitotic rounding just prior to cytokinesis from lumenal pressure and reactionary support from hindbrain tissue (hb, grey). **(B)** Diagram illustrating the deformation of the adjacent hindbrain tissue (hb, grey) as the otic vesicle grows from internal pressure. **(C-F)** 2D confocal micrographs of the otic vesicle at (C) 16 hpf, (D) 24 hpf, (E) 28 hpf, and (F) 32 hpf highlighting the progressive deformation of adjacent hindbrain and ectoderm tissues relative to the dashed-green line. The red and blue arrow heads highlight the progressive deformation in the shape of mitotic cells at contact and non-contact regions. **(G)** Quantification of mitotic cell aspect ratios at contact regions (hindbrain-vesicle or ectoderm-vesicle interface, blue markers) and other non-contact regions (anterioposterior poles, red markers). Aspect ratio is measured as the ratio of apico-basal to lateral cell radii. **(H)** Distribution of division plane orientation relative to the lumenal surface-normal at contact and non-contact cell populations. **(I)** Distribution of division plane orientation for all cells across three stages 16-25, 25-35, and 35-45 hpf respectively. **(J)** Quantification of hindbrain deformation measured as the peak indentation depth (relative to the dashed green line segment in C-F). n=10 embryos per datapoint. Error bars are SD.

To directly determine the presence of pressure within the otic vesicle, we developed a novel pressure probe able to accurately measure small pressures in small volumes of liquid. This probe consists of a solid-state piezo-resistive sensor coupled to a glass capillary needle filled with water (Figure 3A-B, see STAR METHODS). This device is capable of measuring pressure differences of 5 Pascals (≈0.5mm of water depth) across the range of 50-400 Pascals (Figure S2A). Prior to 30 hpf, lumenal pressure is too low for the needle to penetrate the epithelium. From 30 hpf onwards, we observed that the needle can penetrate into the otic vesicle with no observable volume change due to leakage around the needle (Figure 3B’). Pressure is transmitted from the otic vesicle lumen through the needle tip to the sensor. Readings after puncture increased gradually before reaching a stable pressure level (Figure S2B-D). The positive pressure remained until the glass capillary was withdrawn from the otic vesicle, after which the pressure reading dropped to the baseline value (hydrostatic pressure of the buffer due to its depth in the petri dish), further indicating that a pressure difference exists across the membrane. We measured the pressure level at 30, 36 and 48 hpf and found that the pressure level gradually increases from 100 Pa to upwards of 300 Pa (Figure 3C, S2B-D). These values are similar to prior measurements of the much larger inner ear of adult guinea pigs (Feldman et al., 1979).

**Figure 3:**
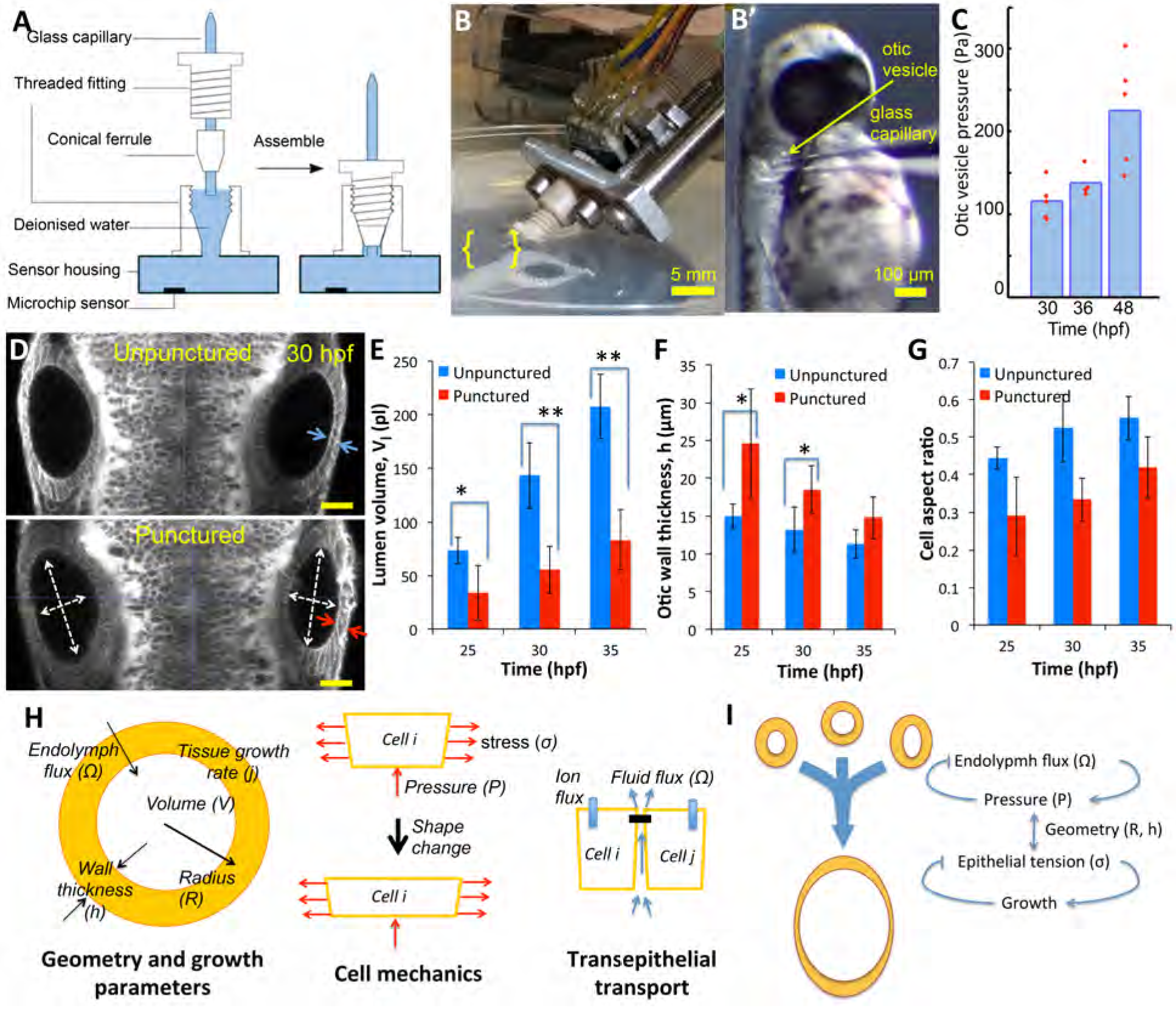
Lumenal pressure drives otic vesicle growth. Pressure measurements in the otic vesicle using a piezo-resistive solid-state sensor. **(A)** The capillary-based probe is mounted on a micromanipulator and zebrafish embryos are immobilized and mounted in Danieau buffer. **(A’)** Under a stereo microscope, the glass capillary is inserted into the otic vesicle. **(B)** Schematic drawing of the pressure probe assembly, not to scale. **(C)** Otic vesicle pressures at different develomental stages of wildtype zebrafish embryos. **(D)** 2D confocal micrographs showing both ears at 30 hpf before (top) and after (bottom) unilateral puncture of the right vesicle. Changes in cell shape from squamous (blue arrows) to columnar (red arrows) are shown. Scale bar is 25 μm. **(E-G)** Quantification of changes from puncturing: (E) lumen volumes (*V*_*l*_, n=10, *p<1.0e-4,**p<1.0e-5), (F) average vesicle wall thickness (*h*, n=10, *p<5.0e-3), and (G) average cell aspect-ratio (n=10, error bars are SD). **(H)** Model relating vesicle geometry, growth rate, and fluid flux to pressure, tissue stress, and cell material properties. **(I)** Multiscale regulatory control of otic vesicle growth linking pressure to fluid transport. Related to Figure S2.

To directly test if lumenal pressure “inflates” the otic vesicle to drive inner ear growth, we punctured otic vesicles at different stages between 25-45 hpf (right vesicle in Figure 3D). Immediately following puncture, we observed a significant decrease in vesicle diameter (white arrows, Figure 3D) and loss of lumenal volume (≈30-40%) (Figure 3E). Examination of punctured vesicles showed that as the vesicle shrunk, the epithelium became thicker (Figure 3D and 3F). Indeed, the excess surface area of the lumenal cavity was absorbed by a significant change in epithelial cell shape to become more columnar while preserving cell volume (in-plane:normal diameter change from 6.7±0.2 μm:13.2±2.9 μm to 5.9±0.2μm:18.4±4.1 μm at 30hpf) (Figure 3G). A similar transition in cell shape is seen when puncturing was conducted at later stages in development (Figure S2K), but importantly to a less columnar resting state suggesting a viscous component. Together, the puncturing experiments provided three insights into the mechanics of the otic vesicle: (i) the lumenal fluid is under hydrostatic pressure that is released when the vesicle is punctured, (ii) lumenal pressure generates stress in the epithelium that alters the shape of epithelial cells, causing them to stretch and become flatter, and (iii) the epithelial tissue response is viscoelastic, being elastic on short time scales, consistent with the epithelium becoming thicker immediately after puncturing, and viscous at longer time scales, consistent with long-term irreversible deformations.

### Theoretical framework linking tissue geometry, fluid flux, and osmotic pressure

Given the complex interplay of lumenal pressure, geometry, and viscoelastic mechanics associated with growth, we sought to develop a mathematical model that accounts for these features (Figure 3H, See STAR METHODS for mathematical model) (Ruiz-Herrero et al., 2017). In a spherically symmetric setting, the relationship between average vesicle radius (*R*), wall thickness (*h*), and tissue growth rate (*j*) can be specified as 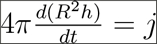. Similarly, the relationship between growth in lumenal volume (*V*_*l*_) and transport across the lumenal surface (of area *S*_*l*_) is related to fluid influx per unit surface area 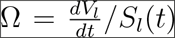. Defining *P*_0_ as the homeostatic pressure required to balance the chemi-osmotic potential driving fluid flux and Ω_0_ as the flux in the absence of a pressure differential, we may write the wildtype fluid flux as Ω = Ω_0_ - *KP*_0_ where *K* is the permeability coefficient (Figure 3F). Intuitively, Ω is the fluid flux that maintains the homeostatic pressure *P*_0_, which in turn, remodels tissue to accommodate the incoming fluid.

Changes in luminal volume can be used to directly determine fluid flux because water is incompressible at low pressures. By using population-averaged measurements of lumenal volume and surface area, we calculate that after an initial rapid expansion (16-20 hpf), flux was approximately constant (Ω ≈ 1*μm*^3^/(*μm*^2^.*hr*) ~ 1*μm*/*hr*) throughout the period 21-45 hpf (Figure 4A). The flux is initially high when there is no pressure but then quickly goes down as pressure builds. Thereafter, fluid accumulation can only occur through viscous expansion of the vesicle. Interestingly, analysis of the system of equations in our model shows that the vesicle will adjust endolymph flux to account for perturbations to vesicle size, via a mechanical feedback loop that links pressure to flux (Figure 3I). Such a control system could be useful to correct natural as well as experimentally induced asymmetry across the left-right axis as we have found in early zebrafish inner ear development (Green et al., 2017).

**Figure 4:**
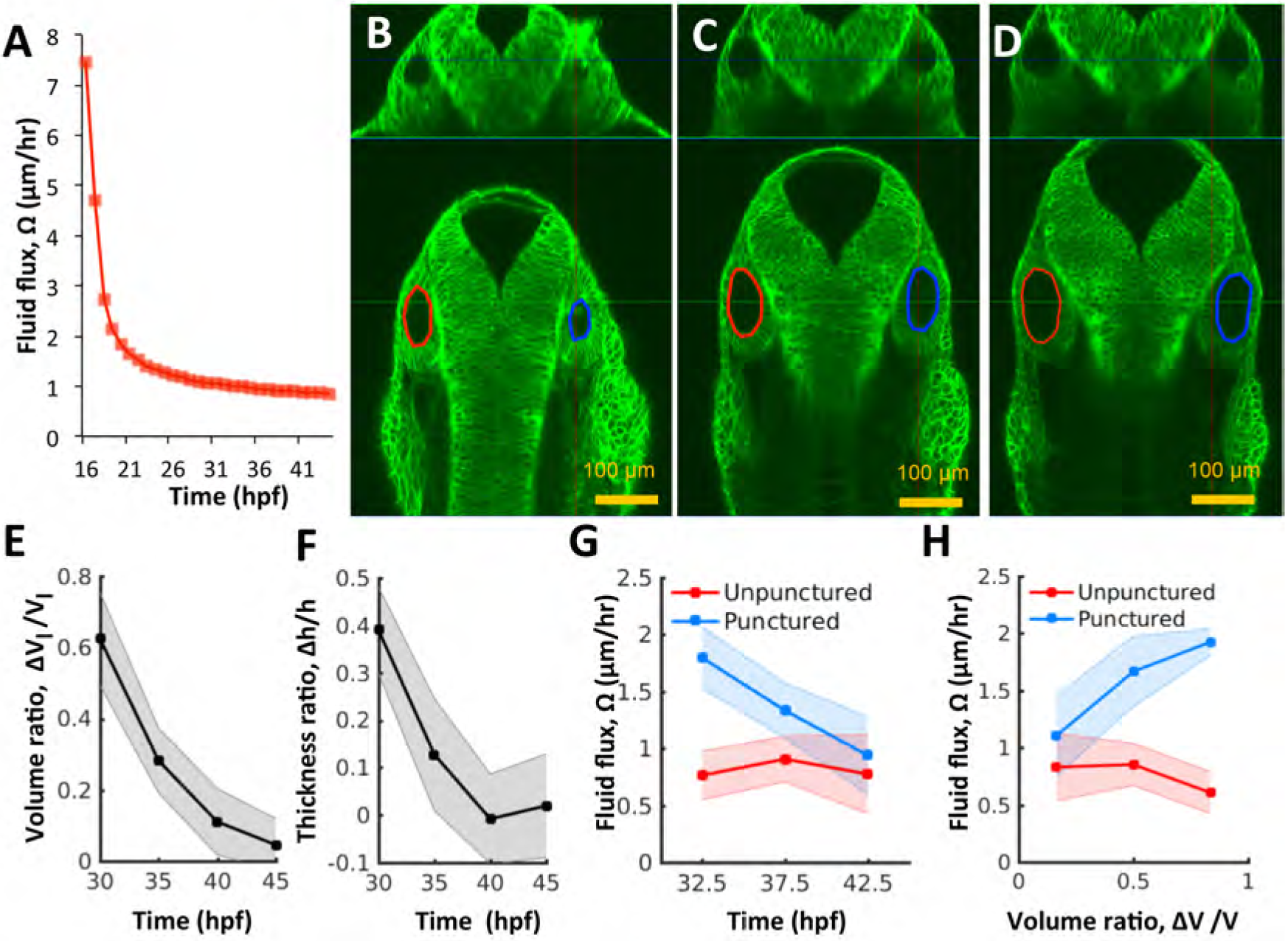
Pressure negatively regulates fluid flux. **(A)** Numerical calculation of fluid flux (Ω) as a function of time using Equation 6 by fitting quadratic polynomials to volume and surface area data. **(B-D)** Confocal 2D micrographs with XZ (top) and XY (bottom) planes depicting the regeneration of a punctured right vesicle (blue) relative to the unpunctured vesicle (left) from (A) 30 hpf right after puncture, to (B) 32.5 hpf, and to (C) 35 hpf. **(E-F)** Quantification of the recovery of volume and wall thickness symmetry. The y-axis plots the difference in lumenal volumes normalized to the unpunctured lumenal volume (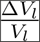, E) and similarly for wall thickness (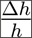, F). Error bars are SD. **(G)** Fluid flux Ω in the punctured ears (blue) and unpunctured ears (red). Error bars are SD. **(H)** Scatterplot showing Ω as a function of volume asymmetry 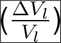 in punctured (blue) and unpunctured (red) ears. Related to Figure S3 and Movie S2.

### Model prediction and validation: Pressure negatively regulates fluid flux

The model predicts that loss of fluid from the lumen (such as by puncture) should lead to a pressure drop and an increased rate of fluid flux back into the lumen and thus a higher than normal growth rate until size is restored, a phenomenon called catch-up growth. To test these model predictions, we experimentally examined whether pressure and fluid flux couple to each other to result in force-based feedback control of development. We first examined the response of the otic epithelium to puncturing perturbations between 25-45 hpf. We injected fluorescent dye (Alexa Fluor 594, 759 MW) into the fluid outside the inner ear (the perilymph) and tracked its movement into the lumen (Figures S2G). Puncturing the otic vesicle and withdrawing the needle created a loss in lumenal volume and allowed the dye from the perilymph to move into the lumen immediately (Figure S2H,I). However, when the dye was injected into the perilymph 5 minutes after the puncture, there was no rapid movement of dye into the lumen (Figure S2J). This showed that the otic epithelium rapidly seals after puncture and restores the epithelial barrier.

Next, we punctured the vesicle at 30 hpf, withdrew the needle, and evaluated its growth relative to the unpunctured contralateral vesicle (control) from 30-45 hpf by simultaneously imaging both otic vesicles. Interestingly, we observed the complete regeneration of lumenal volume in punctured vesicles by an increased growth rate relative to wild type to restore bilateral symmetry (Figure 4B-E, Movie S2). The cell shape changes were also reversed (Figure 4F) suggesting that the vesicle was re-pressurized. The rate of regeneration from fluid flux (Ω) was high immediately after puncture with a slow, gradual decay as bilateral symmetry is restored (Figure 4G). During the rapid recovery phase, Ω in the punctured vesicle (blue curve) was 2-5X higher than that in the unpunctured vesicle (red curve). Our model predicts that upon loss of pressure *P* ≤ *P*_0_ from puncturing, the vesicle dynamically adjusts the fluid flux 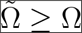 in linear proportion to volume lost. To test this prediction, we pooled data from multiple punctured embryos regenerating from varying levels of pressure loss, to measure how fluid flux related to the volume loss. Consistently, fluid flux in the punctured vesicle correlates with the difference in lumenal volumes between the left and right vesicles (blue markers in Figure 4H and Figure S3 for other developmental stages). These data together show that vesicle pressure negatively regulates fluid flux and suggest that this feedback could buffer variations in size and drive catch-up growth in the otic vesicle.

### Ion pumps are required for lumenal expansion

The transport of salts and fluid across an epithelium can occur through a variety of mechanisms involving transcytosis, electrogenic pumps/transporters and aquaporins for transcellular or paracellular flow (Preston et al., 1992; Hill and Shachar-Hill, 2006; Fischbarg, 2010). Paracellular transport refers to the transfer of fluid across an epithelium by passing through the intercellular space between the cells. This is in contrast to transcellular transport, where fluid travels through the cell, passing through both the apical membrane and basolateral membrane. Previous work in the chick otic vesicle identified the activity of Na^+^-K^+^-ATPase in setting up a transmural potential (Represa et al., 1986) to drive the selective movement of water and ions. In the zebrafish, a role for ion pumps in ear growth is supported by the previous identification of the Na^+^-K^+^-Cl^−^ transporter Slc12a2 as defective in *little ear* mutants (Abbas and Whitfield, 2009). We administered ouabain, an inhibitor of Na^+^-K^+^-ATPase pump activity, to embryos at the 20 hpf stage and quantified vesicle morphology at 30 hpf. We observed a dose-dependent decrease in otic vesicle volume (blue) and wall thickness deformation (red) compared to the wildtype values (Figure 5A) consistent with previous work (Hoijman et al., 2015). In punctured embryos at 25 hpf, adding 500 μM ouabain to the buffer completely inhibited further growth (left vesicle) and post-puncture regeneration (right vesicle in Figure 5B-C, Movie S3). Knockdown of Na^+^-K^+^-ATPase expression in morpholino-injected embryos inhibited lumenal fluid transport in a dose-dependent manner (Figure 5D). We additionally find that otic vesicle growth is sensitive to variations in extracellular pH and blockers of chloride channel activity (Figure 5E-F). Together, these data argue that a network of ion transporters for *Na*^+^, *K*^+^, *H*^+^, and *Cl*^−^ is required for fluid flux into the lumen.

**Figure 5:**
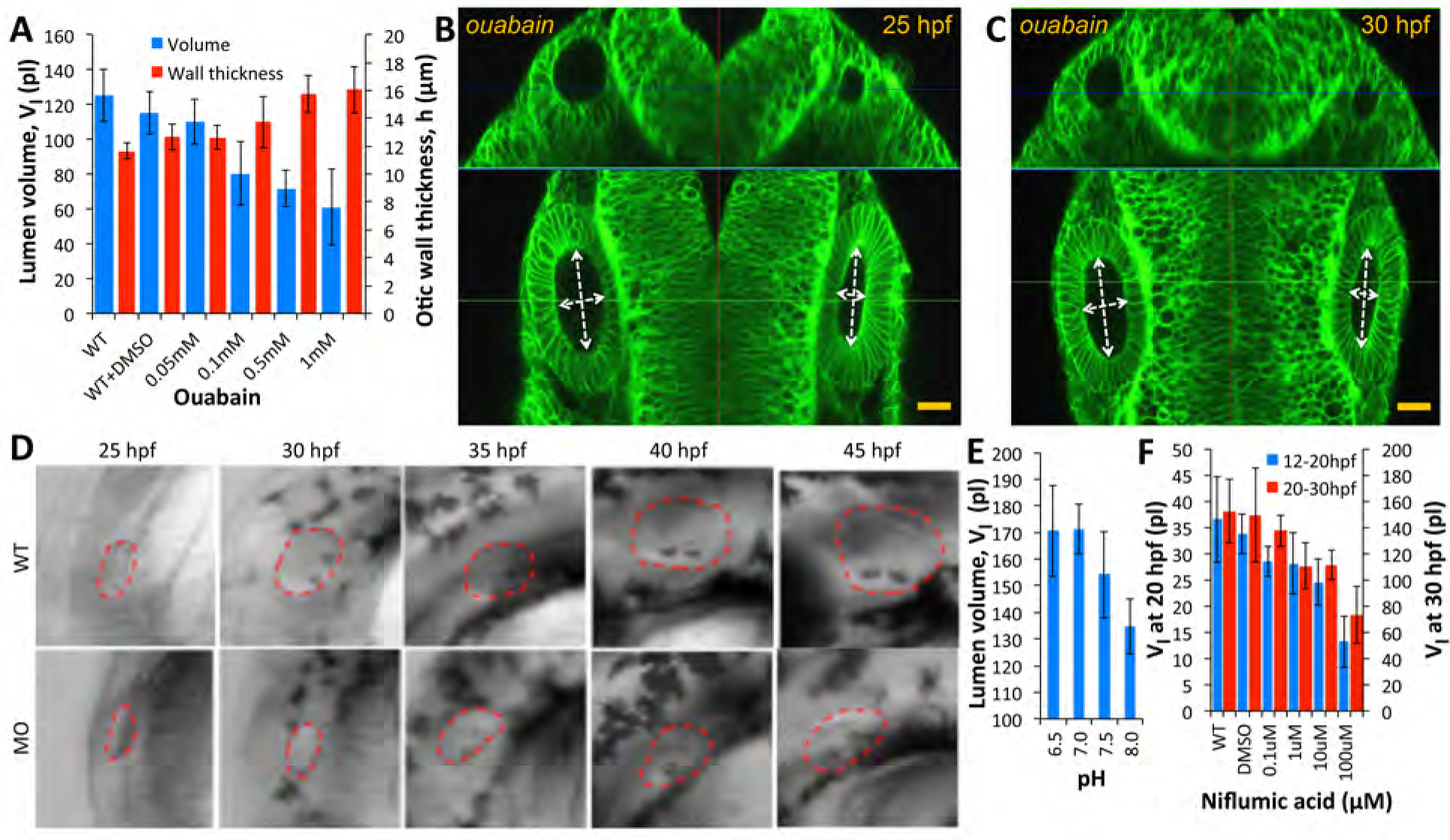
Ear size is affected by disruptions in ion-channel function. **(A)** Quantification of lumenal volume (*V*_*l*_) and wall thickness (*h*) at 30 hpf after ouabain treatment at 20 hpf. Error bars are SD. **(B-C)** Confocal micrographs showing the inhibition of growth in unpunctured (left) and punctured (right) vesicles after incubation in 100μM ouabain to the buffer at 25 hpf. Scale bar 25 μm. **(D)** Brightfield images comparing the growth (25-45hpf) of the wild-type otic vesicle against the antisense morpholino (0.25 ng) targeting the translation of Na, K-ATPase *α*1a.1 mRNA. **(E)** Quantification of lumenal volumes *V*_*l*_ at 30 hpf after acidification of buffer (pH of 6.5-8.0) at 12 hpf. **(F)** Dose-dependent decrease in lumenal volumes *V*_*l*_ observed after the addition of Niflumic acid, a chloride channel inhibitor at 12 hpf (blue) or 20 hpf (red). Related to Movie S3.

### Patterning of tissue material properties causes local differences in epithelial thinning

Our minimal mathematical model assumes that the vesicle is spherical allowing us to understand and predict the qualitative trends of our experiments. For pressure *P* acting inside a thin spherical shell, the tensional tissue-stress—the force pulling cells apart that arises from the radially-outward pushing force of hydrostatic pressure—is 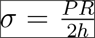. Since the tissue is elastic on short timescales and viscous on long timescales, the radial strain-rate—the change in radius of the otic vesicle, 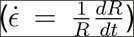—may be related to *σ* via the constitutive relation for a Maxwell fluid given by 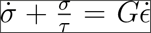, where *G* is the tissue shear modulus—the material property that relates force experienced to deformation—and *τ* is the ratio of the tissue viscosity *μ* and elasticity *k*.

For long timescales, we can use Stokes’ law and force balance (eq. 14, 16, STAR METHODS) to derive an effective tissue viscosity where

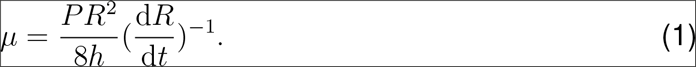

Using this relationship, our morphodynamic measurements, and pressure measurements we estimate the effective viscosity of the otic vesicle tissue to be about 6.4±0.30 × 10^6^ Pa*s from 24-36 hpf and then 2.3±0.13 × 10^7^ Pa*s from 36-45 hpf (see STAR METHODS for error propagation calculations). These values are within the range of tissue viscosities that had been experimentally measured, (Gordon et al., 1972), and indicate that the otic vesicle’s tissue becomes more viscous through development.

Since the vesicle is not actually spherical (Figure S1N,O) and the epithelium is not uniform in thickness (Figure 1H, (Hoijman et al., 2015)), we examined whether (i) the non-spherical vesicle shape creates non-uniform stress distribution as in Laplace’s law and/or (ii) non-uniform patterning of the material properties produce differential strain among cells. To test the first of these possibilities, we tracked anteroposterior pole cells (future sensory) and examined their shapes as they moved from high-curvature to low-curvature regions of the lumenal surface due to epithelial tread milling caused by regional differences in proliferation and emigration (Movie S1). We found that cells retained their columnar shapes independent of the underlying tissue curvature, suggesting that material property patterning may instead contribute to differential cell strain-rates (Figure S4A,B). To test the second possibility, we quantified spatial differences in elasticity (*k*) and viscosity (*μ*) of the otic vesicle using puncturing experiments (Figure 6A). We observed that by eliminating pressure by puncture, medial and lateral cells deformed significantly more compared to the pole cells, indicating that they are softer (smaller *k*) (Figure 6B). Likewise, the resting shapes (post-puncture) of medial and lateral cells were gradually stretched out as the otic vesicle progressed in time through catch-up growth, indicative of their lower viscosity (*μ*, Figure 6C). The observation that the medial and lateral cells become more viscous during development (Figure 6C) agrees with the increased effective viscosity we independently derived from our model using measurements of unperturbed otic vesicle growth. Thus, puncturing perturbations to eliminate pressure forces allowed us to measure spatial differences in cell viscoelasticity.

**Figure 6:**
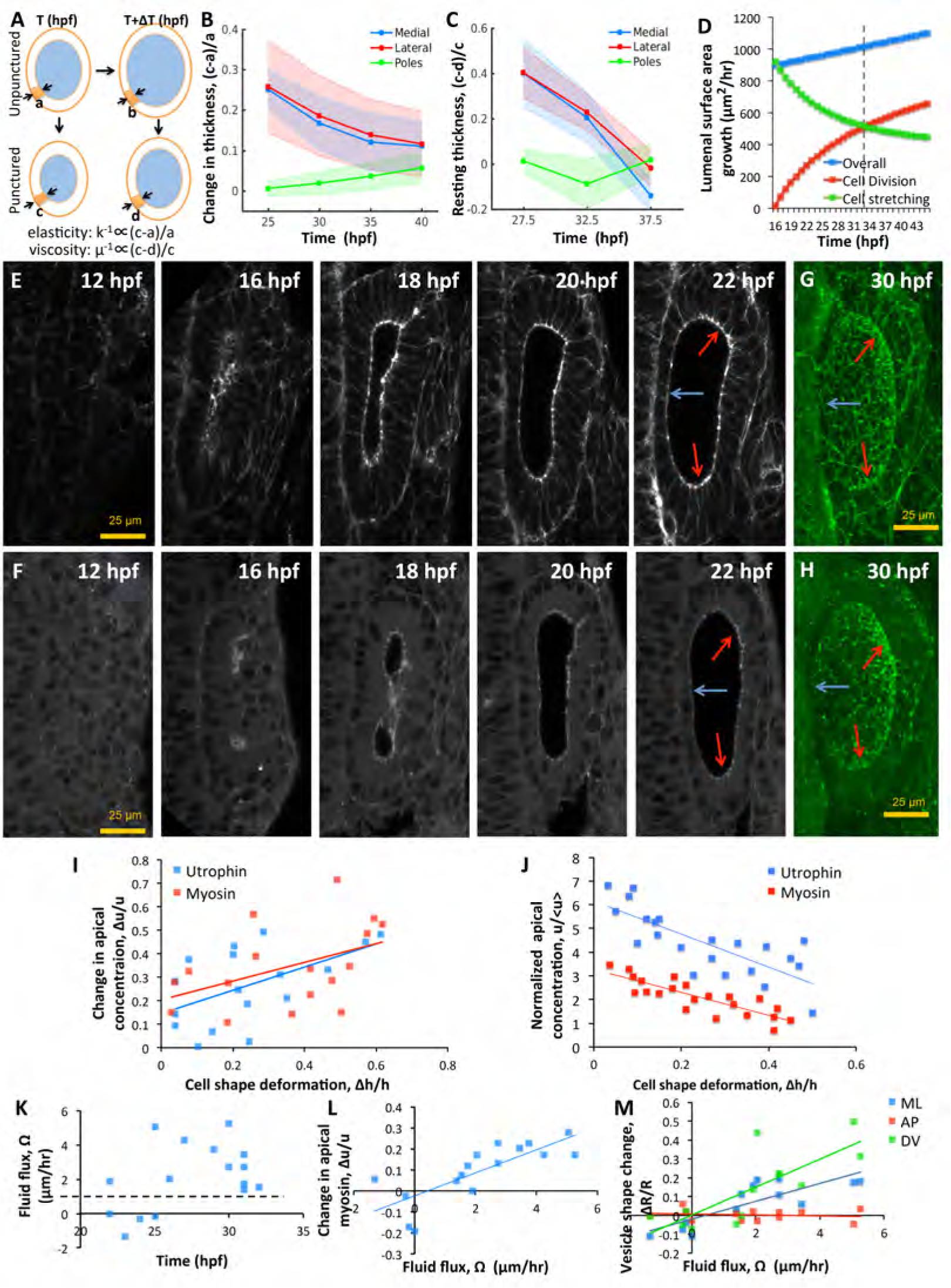
Spatial patterning of material properties results in regional thinning of tissue. **(A)** Schematic for experimentally measuring tissue material properties (*k*, *μ*). The strain of a cell (highlighted in orange) before and after puncturing (*c*−*a*) is inversely proportional to the elasticity modulus *k*. Changes in resting cell-shapes observed over time (*d* − *b*) is inversely proportional to the viscosity parameter *μ*. **(B-C)** Quantification of normalized change in wall thickness ((*c* − *a*)/*a*, B) and *resting* wall thickness ((*c* − *d*)/*c*, C) post-puncture near the hindbrain (medial, blue), ectoderm (lateral, red), and anteroposterior regions (poles, green). Puncturing was done at 25, 30, 35, and 40 hpf. **(D)** Overall lu-menal surface area growth rate (blue markers) showing compensatory contributions from proliferation (red) and cell stretching (green). **(E-F)** Timelapse confocal imaging using *Tg(actb1:GFP-utrCH)* and *Tg(actb1:myl12.1-GFP)* embryos report the dramatic apical localization of F-actin (D) and Myosin II (E) respectively prior to lumenization through 12-16hpf. Through early growth between 16-22 hpf, cells at the poles and lateral regions (red arrows) retain their fluorescence while medial cells lose their fluorescence (blue arrows). **(G-H)** 3D rendering of F-actin (F) and myosin II (G, right) data at 30 hpf show co-localization to apicolateral cell junctions as cells stretch out. **(I)** Quantification of longterm cell shape deformation 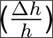 between 16-22 hpf as a function of the rate of change in apical concentration 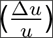 of F-actin (blue markers) and Myosin II (red). **(J)** Quantification of the short-term puncture-induced deformation in cell shapes 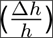 as a function of the normalized apical concentration 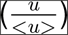 of F-actin (blue markers) and Myosin II (red). < *u* > represents the mean apical expression across the vesicle. Error bars are SD. **(K)** Quantification of fluid flux in embryos treated with 2 mM cytochalasin D at different developmental stages (hpf). Before 25 hpf, embryos failed to grow (Ω ≈ 0) or lose lumenal volume. After 25 hpf, embryos increased their secretion rate by 2-5X over wild-type values (dashed black line, 1 μm/hr). **(L)** Quantification showing the change in apical Myosin II fluorescence 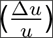 as positively correlated with fluid flux (Ω). **(M)** Quantification of vesicle shape change show maximal change in dorsoventral radius (green markers) compared to the mediolateral (blue) and anteroposterior radius (red). Related to Figure S4 and Movies S4-S7.

### Cell stretching and steady proliferation contribute to tissue viscosity

Our model, calculations, and experimental measurements of cell material properties show that cells progressively become more rigid and more viscous. With diminished ability to remodel cell shape, we examined how otic vesicle growth can be sustained with lumenal pressure. To sustain the same growth rates, our model predicts that overall tissue viscoelasticity should be invariant to changes in cell material properties. While otic tissue elasticity arises from reversible cell stretching (*k*), tissue viscosity is the net result of irreversible cellular stretching (*μ*) as well as proliferation-driven increase in tissue surface area. Thus, we speculated that cell stretching has a more significant role in early growth while proliferation plays a more important role in later stages of growth.

To test these predictions, we evaluated the growth in lumenal surface area (*dS*_*l*_/*dt*) in terms of individual contributions from division 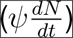 and cell stretching 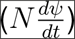 (Figure 6D). Our analysis shows that lumenal surface area growth is linear through time (blue markers). To support this growth, the contribution from cell-stretching is high initially but monotonically decreasing (green markers) and buffered by division (red markers). A breakeven point occurs at around 33 hpf when the contribution to tissue viscosity from cell proliferation exceeds that from cell stretching (dashed black line). Interestingly, our data also show that cell shape stabilizes by this time (Figure 1H). Thus, cell stretching and proliferation play complementary roles through time to sustain a uniform increase in lumenal surface area.

### Tissue material properties are patterned through actomyosin regulation

To identify how cell material properties are patterned, we examined localization patterns of F-actin and Myosin II. Both, F-actin (Figure 6E and Movie S4) and Myosin II (Figure 6F and Movie S5) were apically localized prior to lumenization through 12-16 hpf to form a band around the cavity. Through early growth between 16-22 hpf, gradual and non-uniform changes in the apical density of these molecules are observed. By 30 hpf, we find that these proteins are localized to apicolateral junctions inside cells (Figure 6G-H). Expression levels are retained at pole cells but reduced in medial and lateral cells. Using the movies, we tracked individual cells to understand the relationship between cell shape change and apical expression. We find that wildtype cell deformation during normal growth is positively correlated to localized accumulation of F-actin and Myosin II (Figure 6I). In the transgenic embryos, we used a mosaic labeling strategy for tracking cells to measure the relationship between apical localization and deformation of individual cells immediately following puncture (Figure S4C). We find that upon puncture, the instantaneous deformation observed in individual cells is linearly correlated with the levels of apical localization of F-actin and Myosin II suggesting that actomyosin tension sets effective tissue elasticity (Figure 6J).

To further link spatial patterning of actomyosin localization with epithelial thickness, we conducted loss-of-function experiments and used our model to interpret experimental results. Upon reducing cell elasticity (*k*), our model predicts: (i) an increase in strain-rate 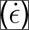 to equilibrate with pressure forces, and (ii) an increase in lumenal dimensions to accomodate increased strain and secretion rates. To decrease cell elasticity, we inhibited actin dynamics by treating embryos at different stages between 16-35 hpf with 100 *μ*M cytochalasin D and used high frame-rate imaging (1 frame/s) to measure vesicle deformations. As predicted by theory, we observed an increase in Ω by a factor of 2-5X over the wildtype values (Figure 6K and Movie S6). The decrease in apical myosin fluorescence positively correlated with the increase in secretion rates (Figure 6L and Movie S6). In these embryos, the DV diameter was found to increase most, compared to LR and AP diameters which experience reactionary forces from the hindbrain and skin (Figure 6M and Movie S6). We also observed that embryos between 16-25 hpf lost volume, presumably, due to a loss in epithelial connectivity and lack of pressure needed for deformation (Movie S7). Together, these data show how spatial patterning of the actomyosin cytokskeleton can lead to spatially varied strain in responses to spatially uniform pressure, and thus contribute to regional differences in the otic vesicle epithelium during growth.

## DISCUSSION

Here, we show that hydrostatic lumenal pressure develops in the zebrafish otic vesicle in response to fluid transport across the otic epithelium to drive growth. We used *in toto* imaging and newly developed quantitative image analysis tools to track changes in cell number, tissue volume, and vesicle lumen volume—which is fluid flux because water is nearly incompressible. We developed a pressure probe device that is amenable to low pressures and small volumes, which enabled us to quantify a developmental increase in hydrostatic pressure. Furthermore, we identified and characterized a new instance of catch-up growth that we leveraged to develop a theoretical framework for otic vesicle size control. With the aid of a multiscale mathematical model, we hypothesized and experimentally confirmed the presence of a hydraulic negative feedback loop between pressure and fluid transport for achieving size control. Modeling helped us systematically integrate the individual contributions of cell physioligical mechanisms underlying pressure and fluid flux, cell proliferation and shape, vesicle geometry, tissue strain-rate and viscoelasticity, to show how growth of the early otic vesicle is controlled. The negative feedback architecture that we found is similar to the chalone model of size control in that the act of growth feeds back to inhibit the rate of growth. However, compared to chalones or morphogen based growth control strategies which are limited in speed by diffusion, the pressure based strategy allows nearly instant communication between different parts of a tissue via hydraulic coupling to allow for “course corrections” to developmental trajectories. Indeed, our study is in line with the fact that hydraulic interactions are relevant to the developmental growth of many internal organs with vesicular and tubular origins (Ruiz-Herrero et al., 2017), and consistent with earlier suggestions that mechanical stresses can play a similar role in regulating organ size (Shraiman, 2005).

### Cause of cell thinning and origin of endolymph

Prior work found that when cells enter into mitosis and round up, their neighbors are stretched and become thinner (Hoijman et al., 2015). This was interpreted as being the mechanism by which cells thin to increase the surface area of the otic vesicle. Here we show that the otic epithelium is fairly elastic and cells can re-thicken following loss of lumenal pressure when the vesicle is punctured (Figure 3F,G), so stretching by neighboring mitotic cells may be short lived. Rather, in our model we postulate that sustained cell thinning during otic vesicle development is caused by in-plane epithelial tension in response to lumenal pressure. It was also noted that cells decrease in volume during early stages of otic vesicle growth and suggested that volume lost from cells is used to inflate the lumen (Hoijman et al., 2015). We attribute the reduction in cell size during early otic development to be due to cell division, and note that the net tissue volume (number of cells multiplied by average cell size) is constant at this stage (Figure 1F). Further, our measurements show that the lumen volume continues to grow until it exceeds the tissue volume (Figure 1F). We thus infer that transepithelial fluid flow is the primary if not sole source of fluid accumulation in the lumen.

### Potential Application of the Lumen Growth Model to Other Systems

Our model for vesicle growth integrates a range of cellular behaviors including division, transport, force generation, material property patterning, and tissue thinning. By tailoring our model equations to different geometries and growth parameters, a unified mathematical framework can be realized to understand size control in hollow organs including the eyes, brain, kidneys, vasculature, and heart. The advantages of such a mesoscale model are several. First, a mesoscale model can be more easily applied to other contexts since the level of abstraction is higher making it less dependent on the specifics of the original context (fewer parameters). Second, growth kinetics and geometry parameters can be experimentally measured using *in toto* imaging approaches. New optical technologies for measuring tissue stresses *in vivo* using oil droplets (Campàs et al., 2014) and laser-ablation (Campinho et al., 2013; Hoijman et al., 2015), ionic gradients using fluorecent ionophores (Adams and Levin, 2012), and pressure (Link et al., 2004) via probes—like the one developed here—hold promise in providing reliable biophysical measurements necessary for understanding morphogenesis. And finally, the contribution of different molecular pathways in regulating model parameters can be prioritized for experimental investigation.

### Comparison of pressure-regulation mechanisms

Our integrated approach combining quantitative imaging and theory-guided experimentation allowed us to identify a novel hydraulic-based mechanism for regulatory control of 3D vesicle growth. This mechanism enables long-range, fast, and uniform transmission of force and connects effects at multiple scales from global pressure forces to supracellular tension and cell stretching mechanics to molecular-scale actions of ion pumps and actomyosin regulation. Later, in the adult ear, tight control of inner ear fluid pressure and ionic composition is necessary to properly detect sound, balance, and body position. Pressure is also important to maintain the structural integrity of organs. Dysregulation of pressure homeostasis can give rise to diseases including hypertension in the vasculature, Meniere’s disease in the inner ear, glaucoma in the eye, and hydrocephalus in the brain. Pressure homeostasis mechanisms may vary. In the inner ear, pressure is initially regulated by feedback between pressure and lumenal fluid flux. However later in development, we have found that a physical pressure relief valve is necessary for pressure homeostasis in the inner ear (Swinburne et al., 2018). Together, we expect that these new insights on ear development and physiology will be critical to the development of effective clinical therapies for hearing and balance disorders, and for understanding size control in closed epithelial tissues.

## AUTHOR CONTRIBUTIONS

K.R.M, I.A.S., A.A.G., N.D.O., and S.G.M. designed the experiments and interpreted the data. C.C. and L.M. designed the pressure probe, K.R.M. and I.A.S. performed imaging experiments. K.R.M, L.M. and S.G.M. developed mathematical models to guide the experiments. K.R.M. and S.T. performed image analysis. K.R.M., I.A.S., L.M. and S.G.M. wrote the paper.

## ACKNOWLEDGEMENTS

We thank members of Megason lab for feedback, Mr. Dante D’India for fish care, and Ms. Suzanne Mosaliganti for help in proof-reading the manuscript. This work was supported by the National Institute of Health grant K25 HD071969 (K.R.M.), Novartis Fellowship for Systems Biology (I.A.S.), National Institute of Health grant 5F32HL097599 (I.A.S.), A*STAR International Fellowship (C.UC.), and Hearing Health Foundation (I.A.S.), the MacArthur Foundation (L.M.), National Science Foundation grant BMMB 15-36616 (L.M.), and National Institute of Health grant R01 DC010791 (S.G.M.).

## STAR METHODS

### CONTACT FOR REAGENT AND RESOURCE SHARING

Further information and requests for resources and reagents should be directed to and will be fulfilled by Lead Contact, Sean Megason.

### EXPERIMENTAL MODEL AND SUBJECT DETAILS

Embryos were collected using natural spawning methods and the time of fertilization was recorded according to the single cell stage of each clutch. Embryos are incubated at 28°C during imaging and all other times except room temperature during injection and dechorionation steps. Staging was recorded using hours post-fertilization as a measure and aligned to the normal table (Kimmel et al., 1995).

#### Zebrafish strains and maintenance

The following fluorescent transgenic strains were used in this study: (i) nuclear-localized tomato and membrane-localized citrine (*Tg(actb2:Hsa.H2B-tdTomato); Tg(actb2:mem-citrine)*^*hm32,33*^), *Tg(actb2:mem-citrine-citrine)*^*hm30*^ (ii) *Tg(actb1:myl12.1-GFP)*^*e2212*^ for visualizing myosin II distribution, and (iii) *Tg(actb1:GFP-utrCH)*^*e116*^ for visualizing F-actin distribution (Behrndt et al., 2012). All fish are housed in fully equipped and regularly maintained and inspected aquarium facilities. All fish related procedures were carried out with the approval of Institutional Animal Care and Use Committee (IACUC) at Harvard University under protocol 04877. Full details of procedures are given in Extended Experimental Procedures.

### METHOD DETAILS

#### Timelapse confocal imaging

A canyon mount was cast in 1% agarose from a Lucite-plexiglass template and filled with 1X Dannieau buffer (Figure S1A). The template created 3 linear-ridges of width 400 μm, depth of 1.5 mm, and length 5 mm (Figure S1B). Canyon-mounted embryos developed normally for at least 3 days with a consistent orientation (lateral or dorsal or dorso-lateral) and can be continuously imaged during this time. Embryos at 15 hpf stage were dechorionated using sharp tweezers (Dumont 55) and mounted dorsally or dorso-laterally (Figure S1C,D) into the immersed canyon mount with a stereoscope (Leica MZ12.5). Multiple embryos for concurrent imaging were mounted in arrays (Figure S1B). Live imaging was performed using a Zeiss 710 confocal microscope (objectives: Plan-Apochromat 20X 1.0 NA, C-Apochromat 40X 1.2 NA) with a home-made heating chamber maintaining 28°C. For experiments requiring the imaging of both left and right ears in an embryo simultaneously, embryos were mounted dorsally and a Plan-Apochromat 20X 1.0 NA objective was used. The inner ear is situated closest to the embryo surface when viewed along the dorso-lateral axis. Dorso-lateral mounting permitted the imaging of the entire ear structure with the best resolution and minimizes the depth of imaging. High-resolution imaging with a Plan-Apochromat 40X 1.2 NA objective facilitated the use of automated image analysis scripts for cell and lumen segmentation, and tracking the movement of fluorescent dyes. Laser wavelengths 488nm, 514nm, 561nm and 594nm lasers were used for confocal time-courses and other single Z-stacks. Embryos were immobilized by injecting 2.3 nl of 20ng/μl *α*-bungarotoxin mRNA (paralytic) at the 1-cell stage for experiments requiring long-term time-lapse imaging (Swinburne et al., 2015).

##### Wildtype growth curves

The process of anaesthetizing an embryo to prevent twitching and preparing an embryo for continuous imaging can alter wildtype growth dynamics in the long-term. Therefore, we collected single Z-stacks for separate sets of embryos (*N* =10-15) to establish the wildtype growth curves at hourly intervals between 16-45 hpf. These sets of embryos were immobilized rapidly by soaking in 1% tricaine.

##### Confocal microscope settings

Image settings vary by brightness of signal from maternal deposit. For example, (please see Movie S1): labels:membrane-citrine; lasers: 514nm (20mW, 3%); objective: lan-Apochromat 40X 1.2 NA at 1.0 zoom; pixel dwell time: 1.58μs; pinhole size: 89μm; line averaging: 1; image spacing: 0.2×0.2 μm, and 1024×1024 pixels per image, with an interval of 1.0 μm through Z for 80 μm and a temporal resolution of 2 min. The starting Z location for the embryo is ≈20 μm above the top of animal pole to allow sufficient space for it to stay in the field of view or sink in the agarose (Figure S1E,F). A total of 25 control time-lapse (covering the developmental time-period of 15-45 hpf), 450 control Z-stacks (covering the developmental time-period of 15-45 hpf), 45 perturbation-related time-lapse datasets, and 65 perturbation-related Z-stacks were collected for the current report. Embryos were screened for their health before imaging.

#### Vesicular fluid pressure probe

Our pressure probe design was inspired by capillary-based pressure sensing techniques (Husken et al., 1978; Tomos and Leigh, 1999). A piezo-resistive solid-state pressure sensor (Honeywell, HSC series) with high resolution (≈2 Pa, 2 kHz) and minimal mechanical deformation (detailed below) was chosen as the sensing element. The sensor was coupled via a high pressure fitting to a ≈2 cm long glass capillary (World Precision Instruments) with a conical tip of 2-20 μm inner diameter (Figure 3A-B). Before they were coupled, both the capillary and sensor were filled with deionized water and the sensor was carefully degassed to ensure the entire interior volume is filled with water. Thus, the fabricated pressure probe had a water-filled dead-end cavity with the only opening at the capillary tip. A detailed fabrication procedure will be available in a separate publication. The digital output from the sensor was sampled by a developer board at 10 Hz (Arduino Uno-R3, and in-house Matlab program). We calibrated our fabricated pressure probe by measuring the hydrostatic pressure at different water depths (Figure S2A) that matched with the sensor calibration provided by the manufacturer.

To measure the lumenal pressure in otic vesicle, zebrafish embryos were immobilized by injecting *α*-bungarotoxin mRNA and dorsolaterally positioned in a canyon mount as before. The pressure probe was mounted on a micro-manipulator typically used for injection. Puncturing was done under a stereo-microscope: the capillary tip was first placed next to the otic vesicle to measure the reference hydrostatic pressure, and then the tip was punctured into the otic vesicle from the lateral direction. A tight sealing was indicated by the vesicle being intact while the capillary tip was inside the vesicle. Examples of pressure profiles are shown in Figure S2B-D. We took the mean pressure value at the plateau, i.e. after the initial pressure rise and before any drop due to leakage, as the fluid pressure inside the vesicle and the data is presented in Figure 3C in the main text. As a negative control, we also punctured in bulk tissue regions such as in the neural tube and measured no pressure rise.

Since the sensor, glass capillary and otic vesicle form a closed volume after puncture, any deformation on the sensing element is compensated by out-flux of the luminal fluid from the vesicle. To verify that the volume change of the piezoresistive element is negligible, and therefore does not significantly reduce the luminal pressure, we calculated the elastic deformation of the sensing membrane and compare against the otic vesicle volume. We disassembled the sensor housing and measured the membrane area to be a square with edge length *L*=850 μm. The membrane thickness, *h*, is estimated to be 5-50 μm (Ruiz et al., 2012). The material is simplified as an isotropic silicon plate with Young’s modulus *E*=163 GPa (Chicot et al., 1996; Hess, 1996; Dolbow and Gosz, 1996) and Poisson’s ratio *ν*=0.27 (Hess, 1996; Gan et al., 1996). The small transverse displacement, *w*, of a thin plate under an uniform transverse hydrostatic pressure, *P*, can be calculated using the Kirchhoff-Love plate theory (Timoshenko and Woinowsky-Krieger, 1959) ∇^2^∇^2^*w* = *P*/*D*, where the bending stiffness *D* = 2*h*^2^*E*/3(1 − *ν*^2^). The boundary condition for the built-in edges satisfies 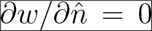, where 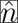 is the in-plane normal of the edges. The transverse deformation field, *w*(*x*, *y*), was obtained by solving the above partial differential equation with a finite element solver (Matlab, Mathwork). An analytical solution exists for the deformation at the center as (Timoshenko and Woinowsky-Krieger, 1959, Article 44) *w*_*c*_ = 1.26 × 10^−3^*PL*^4^/*D*, which agrees well with the numerical solution (Figure S2E). The corresponding volume change, *V* = *∬*_L_ *w*(*x*, *y*)*dxdy*, was depicted in Figure S2F. Comparing to the volume of a 200 μm diameter sphere (dotted line in Figure S2F), the volume change is at least 2 orders of magnitude smaller and hence the resultant pressure drop is negligible.

#### Ouabain and Cytochalasin D treatment

In order to inhibit the activity of Na^+^,K^+^-ATPase, embryos at the 20 hpf stage were soaked in 1% DMSO + ouabain (Sigma Aldrich, CAS 11018-89-6) across a range of concentrations from 0-1 mM. Ouabain was stored at 10 mM in 1% DMSO and diluted to required concentrations in 1X Dannieau buffer before use. Ear size was assessed at 30 hpf as an endpoint. For long-term imaging, ouabain was added to 1% agarose mold used for mounting the embryos. To ensure consistent penetration, 2.3 nl of 0.75 mM ouabain was injected (Nanoject) into the cardiac chamber for circulation throughout the embryo. Assuming an average embryo volume of ~180nl, this injected dose guarantees an effective concentration of 10 μM. In order to perturb the actin network in the otic vesicle, 2.3 nl of 2mM cytochalasin D was injected into the cardiac chamber for an effective concentration of 25 μM in the ear.

#### Buffer pH and Niflumic acid perturbation

To study the effect of pH on otic vesicle size, embryos at the 12 hpf stage were soaked in 1X Dannieau buffer titrated to different pH levels ranging from 6.5-8.5 at 12 hpf. We chose to use the 12 hpf to ensure that the embryo pH homeostasis was adequately perturbed before ear growth commenced. We assessed sizes at 25 and 30 hpf. In order to inhibit the activity of chloride channels and pH regulation in the embryos, embryos at the 20 hpf stage were soaked in 1% DMSO + niflumic acid (Sigma Aldrich, CAS 4394-00-7) across a range of concentrations from 0-1 mM. Ear size and cell shape was assessed at 30 hpf.

#### Antisense morpholino injection

A total of four *α*1-like and two *β* subunit expression of the Na,K-ATPase gene have been identified in the inner ear with distinct spatiotemporal patterns of expression (Blasiole et al., 2003). Antisense morpholino (Gene Tools LLC; Philomath, OR) with sequence (5’- gccttctcctcgtcccattttgctg-3’) targeted against the Na,K-ATPase *α*-subunit gene atp1a1a.1 (*α*1a.1) was developed to knockdown expression in the early otic vesicle (Blasiole et al., 2006). The ability of the morpholino to act specifically to knockdown translation of only the relevant isoform of the Na,K-ATPase mRNA was previously demonstrated using an *in vitro* translation assay (Blasiole et al., 2006). To examine the role of the Na,K-ATPase in controlling ear growth, the morpholino was suspended in 1% DMSO and injected into 1-cell wild-type zebrafish embryos. Here, we report our results from using two different doses consisting of 0.25 ng in Figure 4D. Wild-type embryos injected with 1% DMSO were used as negative controls for assessing ear size at different stages of development. In comparison to wild-type phenotypes, 0.5 ng morphants developed smaller otic vesicles, displayed smaller or absent otoliths, curved tails, and lagged in overall development (data not shown). Higher doses of morpholino injection (> 1 ng) made embryos unhealthy prior to otic vesicle lumenization.

#### Puncturing of otic vesicle

To study the development of pressure in the otic vesicle, embryos were mounted dorsolaterally in a canyon mount (1% agaorse by weight) with 1X Dannieau buffer and an unclipped glass needle was slowly inserted into the otic vesicle. The needle pierced the vesicle in a lateral direction. Puncturing locally affected epithelial connectivity, causing on average 1-2 otic cell and 1-2 ectodermal cell deaths. Lumenal fluid (endolymph) leaked out along the circumference of the needle and the epithelium (Figure S2G,H). Needles were positioned using a micromanipulator. The needle was later slowly withdrawn to study regeneration dynamics. Thereafter, the embryo was re-mounted in a dorsal orientation for imaging both the ears simultaneously.

#### Quantifying the viscoelastic material properties

To identify the viscoelastic material properties of otic cells, we punctured vesicles and noted the deformation in cell shape. The puncturing experiment was carried out at 5-hour intervals (25, 30, 35, 40 hpf) to determine trends in material property patterning (Figure 6A). We used a sample size of *n* = 5 − 10 at each timepoint. The observed deformation in the shape of a cell (before vs. after puncture) located at position *x* and time *t* is inversely proportional to the spring constant (Figure 6B). The rate of change in the resting shapes (after puncture) is inversely proportional to the viscosity coefficient (Figure 6C).

#### Mathematical model: linking geometry, growth, mechanics and regulation

To understand otic vesicle growth, we developed a compact mathematical model that links vesicle geometry, tissue mechanics, and cellular behavior. Quantitative imaging identified aspects of cell behavior including cell division, cell size, cell shape, and material properties as being relevant to the size control problem. The process of realizing a multiscale model enabled the identification of two fundamental mechanisms regulating growth: (i) We identified a negative feedback signal linking the development of hydrostatic pressure to the inhibition of lumenal fluid flux for robust control of size; and, (ii) Spatial patterning of actomyosin contractility affected tissue response to pressure forces to shape the ear. In this section, we elaborate on the development of model equations and explain how theory-guided experimentation allowed us to arrive at the correct representation of the process.

#### Conservation of mass links tissue flux and fluid flux to geometry

Quantitative analysis of vesicle geometry pointed to two critically changing parameters, namely, vesicle radii and tissue wall thickness (Figure S1N,O and Figure 1H). A simple model treats the geometry of the otic vesicle as a spherical shell (Figure 3H) of average radius *R* = (*R*_*o*_ + *R*_*l*_)/2 (μm) and wall thickness *h* = *R*_*o*_ − *R*_*l*_ (μm). As notation, variable subscripts *o*, *l*, and *t* refer to entire otic vesicle, otic lumen, and otic tissue components respectively. To account for changes in geometry parameters from growth, we represent otic tissue growth rate as *j* (*pl*/*hr*) and fluid flux (per unit surface-area) as Ω (μm/hr).

Conservation of tissue mass implies the rate of change of the otic tissue volume *j* = *dV*_*t*_/*dt* is related to the change in tissue geometry. Since the average thickness *h* ≈ 12.27 ± 0.32 μm and the smallest otic vesicle radius *R* ≈ 36.2 ± 0.92 μm at 30 hpf, we can make the assumption that wall-thickness is relatively small compared to the average radius (*h* ≪ *R*), so that

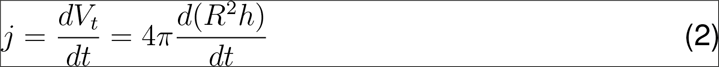

We quantified the parameters *R*, *h*, and *j* and found that average radius increased linearly (Figure S1N,O), wall-thickness (Figure 1H) reduced asymptotically, and tissue volume (Figure 1F) was constant initially (16-28 hpf, *j* = 0) but linearly increased thereafter (28-45 hpf, *j* ≃ 230*pl*/*hr*). Since tissue volume is a product of cell number (*N*) and average cell-size (*s*), we further investigated the role of these cellular parameters during growth.

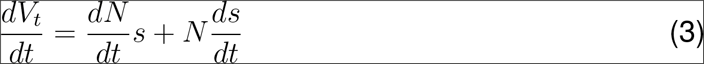

From Figure 1E, cell number *N* monotonically increased non-linearly between 16-45 hpf. In contrast, average cell-size monotonically decreased till 28 hpf and stabilized to a constant value (≃ 0.35*pl*) thereafter. Therefore, between 16-28 hpf, the increase in cell number was offset by a decrease in cell-size (Figure 1E), effectively keeping tissue mass constant. In addition, between 28-45 hpf, increase in tissue mass occurred due to increase in cell number alone with a constant average cell-size.

Similar to volume, we note that lumenal surface area is the product of cell number (*N*) and average cell apical surface area (*ψ*, μm^2^). Therefore, we investigated their individual contribution in driving the increase in surface area.

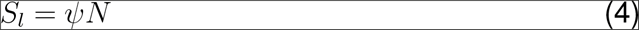

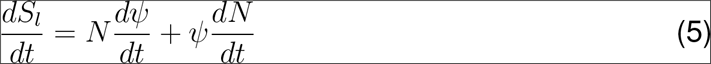

From Figure 1G and 1E, we find that lumenal surface area and cell number increase monotonically from 16-45 hpf. Average apical cell surface showed a saturating response instead. By analyzing both the terms in Figure 6D, we find that both terms contribute significantly to lumenal surface area growth in a temporally complementary manner. Cell stretching 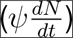 harbors a more significant role during early growth (≤ 32 hpf) with a slow diminishing influence. In contrast, division 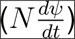 is more significant role during later growth (> 32 hpf). Note that this is a second role of division during growth, in addition to the earlier role in regulating tissue volume increase (Equation 3).

Conservation of fluid volume implies the rate of change in lumenal volume 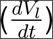 is equal to product of surface area (*S*_*l*_) and fluid flux Ω.

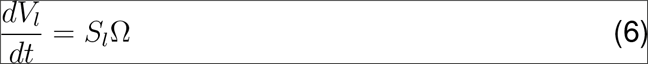

For a spherical geometry, lumenal volume is 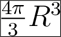 and surface-area is 4π*R*^2^, so that

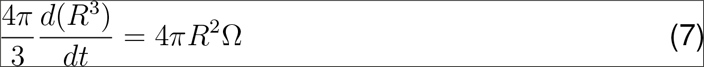

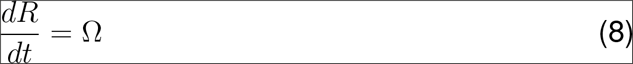

In other words, fluid flux is the same as the rate of change of lumenal radius. From Figure 4A and Figure S1N, we find that average radius increased linearly and lumenal fluid flux was approximately constant (≃ 1*μm*/*hr*) between 16-45 hpf.

Thus, our analysis of growth kinetics using simple conservation laws points to the role of mechanisms involved in regulating cell division, cell volume, fluid transport, and cell shape in controlling size.

#### Modeling pressure generation and feedback to fluid transport mechanisms

Since lumenal volume growth contributes the most to vesicle growth, we examined the phenomenon of fluid transport into a closed cavity (Figure 1F). Prevailing theories of fluid transport suggest the movement of charged ions and fluid through intercellular junctions and channels on cell surfaces (Hill and Shachar-Hill, 2006; Fischbarg, 2010). To model the phenomenon of fluid transport into a closed cavity, we denote the rate of transport of solute as *M*(*mol.μm*^−2^ *.h*^−1^) and its concentration in the lumen as *C* (*mol.μm*^−3^). Then, fluid flux required to retain this concentration is given by the relationship:

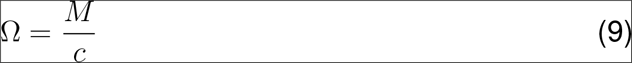

Earlier, our analysis showed that lumenal fluid flux Ω is a constant throughout growth. Under the assumption of isotonic transport (*c* is a constant), we inferred that lumenal solute flux *M* is also constant throughout growth. Since solute and fluid flux are coupled, we investigated which of these parameters are regulated. Two possible scenarios exist: (i) Otic epithelium ensures a solute flux of exactly *M* leading to a fluid flux of 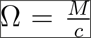, or (ii) Otic epithelium restricts fluid flux to Ω on account of wall distensibility and pressure gradient, which effectively retains a net solute flux of *M* = Ω*c*. Indeed, the existence and presence of lumenal pressure is evident in terms of causing shape deformation in hindbrain tissue and mitotic cells (Figure 2). To concretely show the existence of pressure and how it restricts fluid transport, we performed simple perturbation experiments:

1. Puncturing experiments to verify the presence of pressure by assaying the loss of lumenal volume (Figure 3).
2. Regeneration experiments to demonstrate that the otic vesicle is capable of increased rates of solute and fluid transport in the absence of pressure. We measured regeneration dynamics (Figure 4) and found that vesicles were able to increasing flux by a factor *k* = 3 − 4X the *wild-type* values (Figure S3).

Experimental outcomes suggest that pressure creates an opposing force to the osmotic potential forces to inhibit the movement of fluid, thus setting growth rates appropriate to the developmental stage. Thus, in unperturbed *wildtype* embryos, the observed fluid flux (Ω) is a function of the difference between the osmotically-derived flux (Ω_0_, arising purely due the osmotic potential alone) and the deviation from a set-point pressure, that we term the homeostatic pressure in the vesicle *P*_0_. Making the assumption that the functional form is linear then suggests the equation

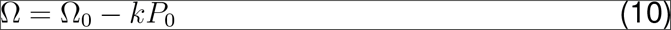

In punctured embryos, the loss in pressure (*P* ≤ *P*_0_) leads to a larger flux following regeneration 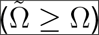.
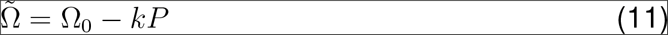

i. e. the increase in flux relative to the unperturbed value is proportional to the loss in pressure relative to its homeostatic value, with

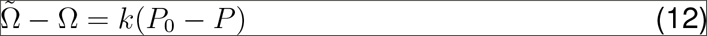

For small changes in the radius in the neighborhood of the fixed point, the loss-in-pressure is proportional to the loss-in-volume (*V*_0_ − *V*), where *V*_0_ and *V* are unperturbed and perturbed volumes respectively), so that we arrive at the following relationship:

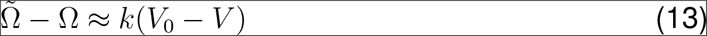

We validate model predictions of a correlation between regeneration flux (Ω) and volume-loss between punctured and unpunctured ears (*V*_0_ − *V*). Our experimental data (regeneration in blue curve in Figure 4H and Figure S3E,F) depicts a proportional relationship, thus validating the accuracy of our model. Our data shows that the unperturbed flux Ω ≃ 1*μm*/*hr* and slope *k* is approximately 3-5.

#### Modeling pressure-driven growth and deformation of the vesicle shape

The otic vesicle is expected to deform under the action of pressure, thus leading to growth and reshaping of the tissue. We therefore sought to assess how these forces affect individual cells. Given the vesicle geometry, cells in the otic vesicle experience three types of pressure-derived forces: (i) pressure force (*P*, *N.μm*^−2^) normal to the apical membranes in a direction that flatten cells, (ii) tissue stress (*σ*, *N.μm*^-2^) distributed normal to lateral membranes, and (iii) reactionary or support forces from hindbrain and skin tissue.

To formalize this, we modify a recent framework introduced to study the growth of cellular cysts (Ruiz-Herrero et al., 2017). In a closed geometry, tissue stress *σ* is related to lumenal pressure *P* by a simple force balance equation. For a spherical pressure vessel, the pressure force pushing one hemisphere is counterbalanced by tissue tension.

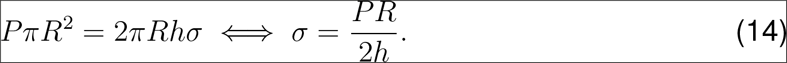

To link tissue stress to deformation, we investigated the material properties of the tissue (*G*). We initially modeled otic tissue as being similar to a purely viscous fluid flowing under a tangential shear stress *σ* with a strain-rate of 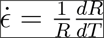.

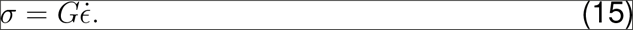

For a purely viscous fluid with viscosity coefficient *μ* (*Pa.h*) deforming in a spherically-restricted geometry, the shear stress (*σ*) in the fluid layers relates to rate of radius change using Stokes’ law as:

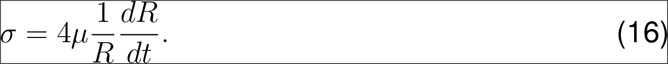

Since a viscous material deforms continuously under the action of a force and retains the deformation when force is removed, we validated our assumption by studying the temporal trends in resting shapes of cells after pressure forces are eliminated. Indeed, we observed that the resting shape of cells (after puncture) had undergone irreversible deformation in developmental time (Figure 3E and 3C (red bars progressively decrease)). In other words, a cell at 35 hpf has a relatively squamous morphology at rest compared to the same cell at 25 hpf. When compared to the contralateral unpunctured ear, however, we observed an elastic response in punctured vesicles wherein cells had dynamically reshaped to more columnar morphologies (Figure 3A and 3C,D (difference between red and blue bars)). This suggested that the tissue behaved elastically in short time-scales and viscous in long time-scales. To incorporate elastic behavior on short timescales, we modeled the overall deformation as:

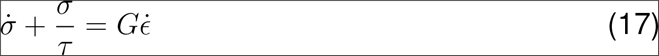

We next quantified cell material properties by quantifying the short (elastic) and long timescale (viscous) responses in response to puncturing perturbations (Figure 6A). The reversible component of a cell’s deformation, when all acting forces are eliminated, is inversely proportional to its elasticity (*k*). The irreversible component of a cell’s deformation that accumulates over time is inversely proportional to the viscosity coefficient (*μ*). We also investigated and identified the molecular origins of material property patterning in terms of actomyosin networks that were found to be apically expressed in the otic cells (Figure 6). Thus, our model identifies the role of pressure forces in deforming cells and driving growth. Material property patterning was found to locally modulate the deformation to anisotropically shape the vesicle through growth.

## QUANTIFICATION AND STATISTICAL ANALYSIS

Each data point of morphodynamic quantification was obtained using our automated bioimage informatics pipeline (ITIAT) on images from 10 different embryos that were immobilized with *α*-bungarotoxin mRNA. 30 time-points were analyzed—every hour between 16 and 45 hpf—which included 300 otic vesicle lumen measurements and more than 200,000 segmented cells. For Figure 1E-H, S1N-O, 2J, the translucent spread of the data is the standard deviation.

For analysis of mitotic cell deformation, we identified 54 mitotic cells. The spread of the data presented as translucent overlays in Figure 2G is the calculated standard deviation. For measuring the hindbrain deformation in Figure 2J, n=10 embryos/timepoint were used.

Pressure measurements were acquired from 5 different embryos at 30, 36, and 48 hpf (Figure 3C).

For puncture experiments, 10 otic vesicles were measured that were punctured and there morphometrics were compared to 10 unpunctured otic vesicles (Figure 3D-F, 4E-H, S3). The error bars are standard deviation and the p-values were obtained using a student t-test.

For ion-channel and pH perturbations, each data point is 5 embryos and error bars are standard deviation (Figure 5).

For calculating the effective tissue viscosityfrom Equation 1, the time-averaged data of all variables were obtained from the datasets shown in Figure 3C,E-F. The instantaneous growth rate, *R*’ = d*R*/d*t*, was calculated from the difference of radius R of each time point before averaging. A formula for error propagation (Ku, 1966) was derived from Equation 1:

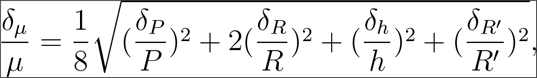

where *δ*_*X*_ is the standard deviation of variable *X*. The mean and standard deviations are summarized in the table below:

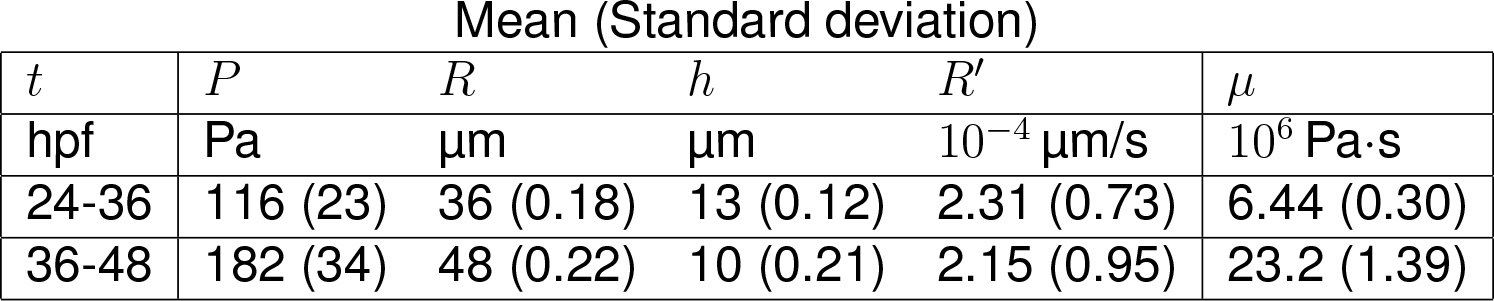

| Mean (Standard deviation) |  |  |  |  |  |
| --- | --- | --- | --- | --- | --- |
| $t$ | $P$ | $R$ | $h$ | $R'$ | $\mu$ |
| hpf | Pa | $\mu\text{m}$ | $\mu\text{m}$ | $10^{-4} \mu\text{m/s}$ | $10^6 \text{ Pa}\cdot\text{s}$ |
| 24-36 | 116 (23) | 36 (0.18) | 13 (0.12) | 2.31 (0.73) | 6.44 (0.30) |
| 36-48 | 182 (34) | 48 (0.22) | 10 (0.21) | 2.15 (0.95) | 23.2 (1.39) |

For the derivation of relative differences in tissue elasticity and viscosity, each data point was obtained from 10 different punctured embryos (Figure 6A-C) and the translucent spread represents the standard deviation.

Figure 6I-J, 22 utrophin expressing cells and 22 myosin expressing cells were analyzed. Figure 6K-L-M, scatterplots with 15, 16, and 12 embryos were used respectively.

## DATA AND SOFTWARE AVAILABILITY

### Image analysis

Our entire bioimage informatics pipeline called *“In toto image analysis toolkit (ITIAT)”* is available online at: https://wiki.med.harvard.edu/SysBio/Megason/GoFigureImageAnalysis. The pipeline can be used for generating 3D surface reconstructions, automatic whole-cell and nuclei segmentations, and cell tracking. The code is open-source and written using the C++ programming language. The code can be downloaded and compiled in any platform using CMake, a tool for generating native Makefiles. The code is built by linking to pre-compiled open-source libraries, namely, ITK (http://www.itk.org) and VTK toolkit (http://www.vtk.org) for image analysis and visualization respectively. We also used the open-source and cross-platform GoFigure2 application software for the visualization, interaction, and semi-automated analysis of 3*D* + *t* image data (http://www.gofigure2.org) (Gelas, Mosaliganti, and Megason et al., in preparation). Measurement of mitotic cell aspect-ratios was carried out using ZEN (Carl Zeiss) software 3D distance functionality. Measurements were analysed and plotted with Matlab (Mathworks) and Microsoft Excel. To obtain 3D models of the otic vesicles, 2D contours were first placed along regularly-sampled z-planes in GoFigure2 (Figure 1I). 3D reconstructions were obtained using the Powercrust reconstruction algorithm (https://github.com/krm15/Powercrust) (Amenta et al., 2001). Automatic cell and lumen-segmentation were performed using ACME software for whole-cell segmentations (Mosaliganti et al., 2012; Xiong et al., 2013) (Figure S1J-M).

### KEY RESOURCES TABLE

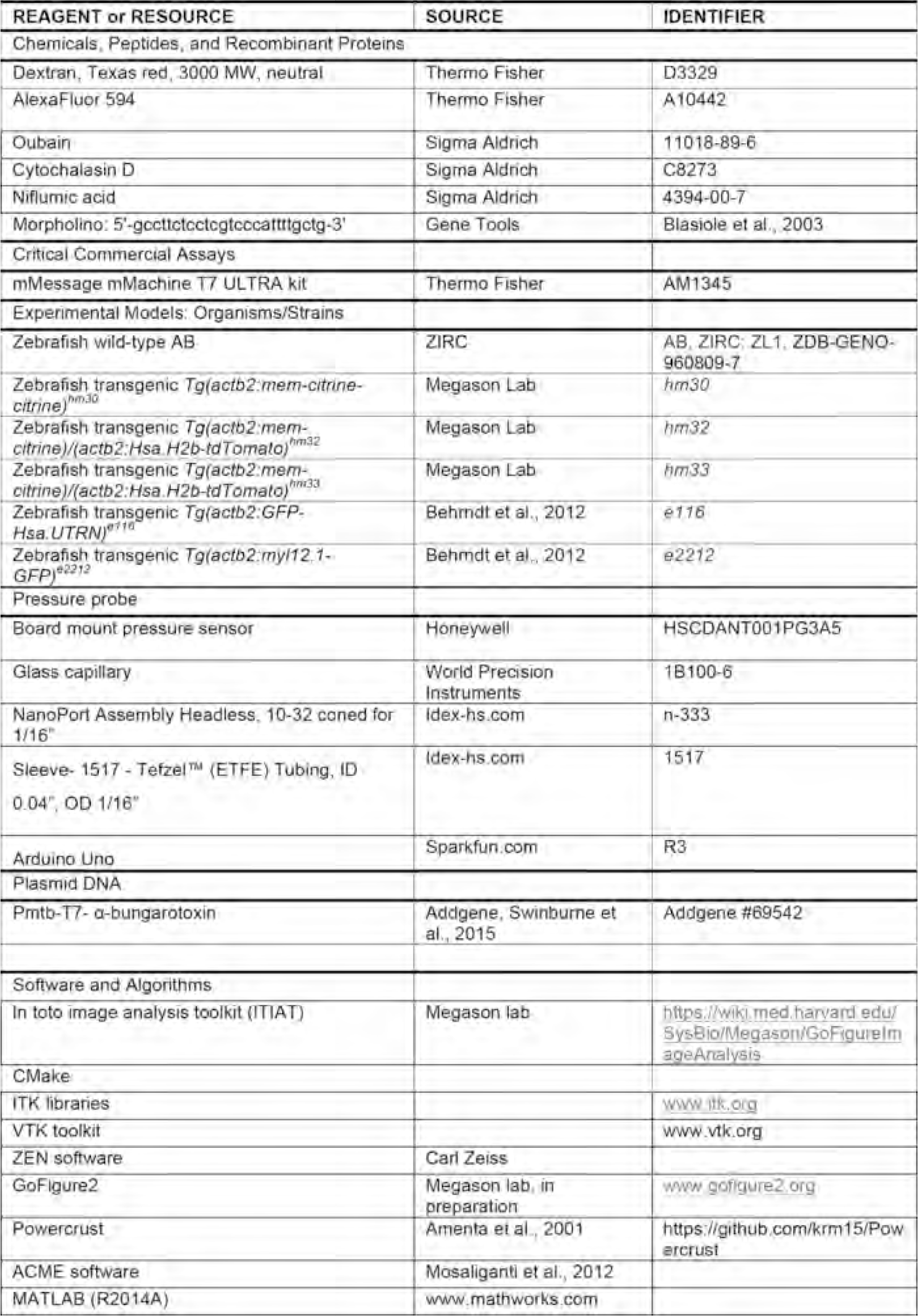

### SUPPLEMENTAL INFORMATION

Supplemental information includes four figures and seven movies.

## REFERENCES

Abbas, L. and Whitfield, T. T. (2009). Nkcc1 (Slc12a2) is required for the regulation of endolymph volume in the otic vesicle and swim bladder volume in the zebrafish larva. Development 136, 2837–2848.

Adams, D. S. and Levin, M. (2012). General principles for measuring resting membrane potential and ion concentration using fluorescent bioelectricity reporters. Cold Spring Harbor Protocols 7, 385–397.

Amenta, N., Choi, S. and Kolluri, R. K. (2001). The power crust, unions of balls, and the medial axis transform. Computational Geometry: Theory and Applications 19, 127–153.

Bagnat, M., Cheung, I. D., Mostov, K. E. and Stainier, D. Y. R. (2007). Genetic control of single lumen formation in the zebrafish gut. Nature cell biology 9, 954–960.

Behrndt, M., Salbreux, G., Campinho, P., Hauschild, R., Oswald, F., Roensch, J., Grill, S. W. and Heisenberg, C.-P. (2012). Forces Driving Epithelial Spreading in Zebrafish Gastrulation. Science 338, 257–260.

Blasiole, B., Canfield, V. A., Vollrath, M. A., Huss, D., Mohideen, M.-A. P. K., Dickman, J. D., Cheng, K. C., Fekete, D. M. and Levenson, R. (2006). Separate Na, K-ATPase genes are required for otolith formation and semicircular canal development in zebrafish. Developmental Biology 294, 148–160.

Blasiole, B., Degrave, A., Canfield, V., Boehmler, W., Thisse, C., Thisse, B., Mohideen, M.-A. P. K. and Levenson, R. (2003). Differential expression of Na, K-ATPase alpha and beta subunit genes in the developing zebrafish inner ear. Developmental Dynamics 228, 386–392.

Bryant, P. and Levinson, P. (1985). Intrinsic growth control in the imaginal primordia of Drosophila, and the autonomous action of a lethal mutation causing overgrowth. Developmental Biology 107, 355–63.

Bullough, W. S. and Laurence, E. B. (1964). Mitotic control by internal secretion: the role fo the chalone-adrenalin complex. Experimental Cell Research 33, 176–194.

Campàs, O., Mammoto, T., Hasso, S., Sperling, R. a., O’Connell, D., Bischof, A. G., Maas, R., Weitz, D. a., Mahadevan, L. and Ingber, D. E. (2014). Quantifying cell-generated mechanical forces within living embryonic tissues. Nature methods 11, 183–9.

Campinho, P., Behrndt, M., Ranft, J., Risler, T., Minc, N. and Heisenberg, C.-P (2013). Tension-oriented cell divisions limit anisotropic tissue tension in epithelial spreading during zebrafish epiboly. Nature Cell Biology 15, 1405–14.

Chicot, D., Dméearécaux, P. and Lesage, J. (1996). Apparent interface toughness of substrate and coating couples from indentation tests. Thin Solid Films 283, 151–157.

Colombani, J., Raisin, S., Pantalacci, S., Radimerski, T., Montagne, J. and Lopold, P. (2003). A Nutrient Sensor Mechanism Controls Drosophila Growth. Cell 114, 739–749.

Coulombre, A. J. (1956). The role of intraocular pressure in the development of the chick eye. Journal of Experimental Zoology 133, 211–255.

Das, D., Chatti, C., Emonet, T. and Holley, S. A. (2017). Patterned disordered cell motion ensures vertebral column symmetry. Developmental Cell 42, 170–180.

Dasgupta, A., Merkel, M., Clark, M. J., Jacob, A. E., Dawson, J. E., Manning, M. L. and Amack, J. D. (2018). Cell volume changes contribute to epithelial morphognesis in zebrafish Kupffer’s vesicle. eLife 7, e30963.

Day, S. J. and Lawrence, P. A. (2000). Measuring dimensions: the regulation of size and shape. Development 127, 2977–2987.

Debat, V. and Peronnet, F. (2013). Asymmetric flies: the control of developmental noise in Drosophila. Fly 7, 70–77.

Desmond, M. E. and Jacobson, A. G. (1977). Embryonic brain enlargement requires cerebrospinal fluid pressure. Developmental Biology 57, 188–198.

Dittmer, J. E., Goss, R. J. and Dinsmore, C. E. (1974). The growth of infant hearts grafted young and adult rats. American Journal of Anatomy 141, 155–160.

Dolbow, J. and Gosz, M. (1996). Effect of out-of-plane properties of a polyimide film on the stress fields in microelectronic structures. Mechanics of materials 23, 311–321.

Dupont, S., Morsut, L., Aragona, M., Enzo, E., Giulitti, S., Cordenonsi, M., Zanconato, F., Digabel, J. L., Forcato, M., Bicciato, S., Elvassore, N. and Piccolo, S. (2011). Role of YAP/TAZ in mechanotransduction. Nature 474, 179–183.

Dyballa, S., Savy, T., Germann, P., Mikula, K., Remesikova, M., pir, R. and Pujades, C. (2017). Distribution of neurosensory progenitor pools during inner ear morphogenesis unveiled by cell lineage reconstruction. eLife 6, e22268.

Feldman, A. M., Bittner, H. R., and Brusilow, S. W. (1979). Measurement of the hydrostatic pressures of the cochlear compartments. Neurological Research 1.

Fischbarg, J. (2010). Fluid transport across leaky epithelia: central role of the tight junction and supporting role of aquaporins. Physiological Reviews 90, 1271–1290.

Frömter, E. and Diamond, J. (1972). Route of passive ion permeation in epithelia. Nature New Biology 235, 9–13.

Gamer, L. W., Cox, K., Carlo, J. M. and Rosen, V. (2009). Overexpression of BMP3 in the developing skeleton alters endochondral bone formation resulting in spontaneous rib fractures. Developmental Dynamics 238, 2374–2381.

Gan, L., Ben-Nissan, B. and Ben-David, A. (1996). Modelling and finite element analysis of ultra-microhardness indentation of thin films. Thin Solid Films 290, 362–366.

García-Bellido, A. (2009). The cellular and genetic bases of organ size and shape in Drosophila. International Journal of Developmental Biology 53, 1291–1303.

Gordon, R., Goel, N. S., Steinberg, M. S. and Wiseman, L. L. (1972). A rheological mechanism sufficient to explain the kinetics of cell sorting. Journal of Theoretical Biology 37, 43–73.

Green, A. A., Mosaliganti, K. R., Swinburne, I. A., Obholzer, N. D. and Megason, S. G. (2017). Recovery of shape and size in a developing organ pair. Developmental Dynamics 246.

Günzel, D. and Yu, A. S. L. (2013). Claudins and the modulation of tight junction permeability. Physiological reviews 93, 525–69.

Haddon, C. and Lewis, J. (1996). Early Ear Development in the Embryo of the Zebrafish, Danio rerio. Journal of Comparative Neurology 365, 113–125.

Hariharan, I. K. (2015). Organ size control: lessons from Drosophila. Developmental Cell 34, 255–65.

Hess, P. (1996). Laser diagnostics of mechanical and elastic properties of silicon and carbon films. Applied surface science 106, 429–437.

Hill, A. E. and Shachar-Hill, B. (2006). A new approach to epithelial isotonic fluid transport: an osmosensor feedback model. Journal of Membrane Biology 210, 77–90.

Hoijman, E., Rubbini, D., Colombelli, J. and Alsina, B. (2015). Mitotic cell rounding and epithelial thinning regulate lumen growth and shape. Nature Communications 6.

Hufnagel, L., Teleman, A., Rouault, H., Cohen, S. and Shraiman, B. (2007). On the mechanism of wing size determination in fly development. PNAS 104, 3835–3840.

Husken, D., Steudle, E. and Zimmermann, U. (1978). Pressure Probe Technique for Measuring Water Relations of Cells in Higher Plants. Plant Physiology 61, 158–163.

Ingber, D. E. (2005). Mechanical control of tissue growth: function follows form. PNAS 102, 11571–11572.

Irvine, K. D. and Shraiman, B. I. (2017). Mechanical control of growth: ideas, facts and challenges. Development 144, 4238–4248.

Kimmel, C. B., Ballard, W. W., Kimmel, S. R., Ullmann, B. and Schilling, T. F. (1995). Stages of embryonic development of the zebrafish. Developmental Dynamics 203, 253–310.

Ku, H. H. (1966). Notes on the use of propagation of error formulas. Journal of Research of the National Bureau of Standards 70.

Kunche, S., Yan, H., Calof, A. L., Lowengrub, J. S., and Lander, A. D. (2016). Feedback, lineages, and self-organizing morphogenesis. PLOS Computational Biology 12.

Legoff, L., Rouault, H. and Lecuit, T. (2013). A global pattern of mechanical stress polarizes cell divisions and cell shape in the growing Drosophila wing disc. Development 140, 4051–9.

Lestas, I., Vinnicombe, G. and Paulsson, J. (2010). Fundamental limits on the suppression of molecular fluctuations. Nature 467, 174–178.

Link, B. A., Gray, M. P., Smith, R. S. and John, S. W. M. (2004). Intraocular pressure in zebrafish: Comparison of inbred strains and identification of a reduced melanin mutant with raised IOP. Investigative Ophthalmology and Visual Science 45, 4415–4422.

Lowery, L. A. and Sive, H. (2005). Initial formation of zebrafish brain ventricles occurs independently of circulation and requires the nagie oko and snakehead/atp1a1a.1 gene products. Development (Cambridge, England) 132, 2057–2067.

McPherron, A. C., Lawler, A. M., and Lee, S. J. (1997). Regulation of skeletal muscle mass in mice by TGF-beta superfamily member. Nature 387, 83–90.

Megason, S. G. (2009). In toto imaging of embryogenesis with confocal time-lapse microscopy. Methods In Molecular Biology Clifton Nj 546, 317–332.

Metcalf, D. (1963). The Autonomous Behaviour of Normal Thymus Grafts. The Australian journal of experimental biology and medical science 41, 437–447.

Metcalf, D. (1964). Restricted Growth Capacity of Multiple Spleen Grafts. Transplantation 2, 387–392.

Mosaliganti, K. R., Noche, R. R., Xiong, F., Swinburne, I. A. and Megason, S. G. (2012). ACME: Automated Cell Morphology Extractor for Comprehensive Reconstruction of Cell Membranes. PLoS Computational Biology 8.

Navis, A., Marjoram, L. and Bagnat, M. (2013). Cftr controls lumen expansion and function of Kupffer’s vesicle in zebrafish. Development 140, 1703–1712.

Nelson, C. M., Gleghorn, J. P., Pang, M.-F., Jaslove, J. M., Goodwin, K., Varner, V. D., Miller, E., Radisky, D. C. and Stone, H. A. (2017). Microfluidic chest cavities reveal that transmural pressure controls the rate of lung development. Development 144, 4328–4335.

Pan, Y., Heemskerk, I., Ibar, C., Shraiman, B. I. and Irvine, K. D. (2016). Differential growth triggers mechanical feedback that elevates Hippo signaling. PNAS 113, E6973–E6983.

Parker, N. F. and Shingleton, A. W. (2011). The coordination of growth among Drosophila organs in response to localized growth-perturbation. Developmental Biology 357, 318–325.

Plikus, M. V., Mayer, J. A., de la Cruz, D., Baker, R. E., Maini, P. K., Maxson, R., and Chuong, C. M. (2008). Cyclic dermal BMP signalling regulates stem cell activation during hair regeneration. Nature 451, 340–344.

Preston, G. M., Carroll, T. P., Guggino, W. B. and Agre, P. (1992). Appearance of water channels in Xenopus oocytes expressing red cell CHIP 28 protein. Science 256, 385–387.

Rao, C. V., Wolf, D. M. and Arkin, A. P. (2002). Control, exploitation and tolerance of intracellular noise. Nature 420, 231–237.

Represa, J. J., Barbosa, E. and Giraldez, F. (1986). Electrical properties of the otic vesicle epithelium in the chick embryo. Journal of Embryology and Experimental Morphology 97, 125–139.

Rogulja, D. and Irvine, K. D. (2005). Regulation of cell proliferation by a morphogen gradient. Cell 123, 449–461.

Rosello-Diez, A. and Joyner, A. L. (2015). Regulation of long bone growth in vertebrates; it is time to catch up. Endocrine Reviews 36, 646–680.

Rosello-Diez, A., Stephen, D. and Joyner, A. L. (2017). Altered paracrine signaling from the injure knee joint impairs postnatal long bone growth. eLife 6, e27210.

Rubashkin, A., Iserovich, P., Hernandez, J. A. and Fischbarg, J. (2005). Epithelial fluid transport: protruding macromolecules and space charges can bring about electro-osmotic coupling at the tight junctions. The Journal of membrane biology 208, 251–263.

Ruiz, P. G., Meyer, K. D. and Witvrouw, A. (2012). Poly-SiGe for MEMS-aboveCMOS Sensors. Springer.

Ruiz-Herrero, T., Allesandri, K., Gurchenkov, B. V., Nassoy, P. and Mahadevan, L. (2017). Organ size control via hydraulically gated oscillations. Development 144, 4422–4427.

Savin, T., Kurpios, N. A., Shyer, A. E., Florescu, P., Liang, H., Mahadevan, L. and Tabin, C. J. (2011). On the growth and form of the gut. Nature 476, 57–62.

Shraiman, B. I. (2005). Mechanical feedback as a possible regulator of tissue growth. PNAS 102, 3318–3323.

Silber, S. J. (1976). Growth of baby kidneys transplanted into adults. Archives of Surgery 111, 75–77.

Stewart, M. P., Helenius, J., Toyoda, Y., Ramanathan, S. P., Muller, D. J. and Hyman, A. A. (2011). Hydrostatic pressure and the actomyosin cortex drive mitotic cell rounding. Nature 469, 226–230.

Swinburne, I. A., Mosaliganti, K. R., Green, A. A. and Megason, S. G. (2015). Improved Long-Term Imaging of Embryos with Genetically Encoded-Bungarotoxin. PLoS ONE 10, e0134005.

Swinburne, I. A., Mosaliganti, K. R., Upadhyayula, S., Liu, T.-L., Hildebrand, D. G. C., Tsai, T. Y. C., Chen, A., Al-Obeidi, E., Fass, A. K., Malhotra, S., Engert, F., Lichtman, J. W., Kirchausen, T., Betzig, E. and Megason, S. G. (2018). Lamellar projections in the endolymphatic sac act as a relief valve to regulate inner ear pressure. eLife in press.

Timoshenko, S. P. and Woinowsky-Krieger, S. (1959). Theory of plates and shells, 2nd ed. McGraw-hill.

Tomos, A. D. and Leigh, R. A. (1999). The Pressure Probe: A Versatile Tool in Plant Cell Physiology. Annual Review of Plant Physiology and Plant Molecular Biology 50, 447–472.

Tumaneng, K., Russell, R. C. and Guan, K. L. (2012). Organ Size Control by Hippo and TOR Pathways. Current Biology 22, R368–79.

Twitty, V. C. and Schwind, J. L. (1931). The growth of eyes and limbs transplanted heteroplastically between two species of Amblystoma. Journal of Experimental Zoology 59, 61–86.

Waddington, C. H. (1959). Canalization of development and genetic assimilation of acquired characters. Nature 183, 1654–1655.

Wartlick, O., Mumcu, P., Kicheva, A., Bittig, T., Seum, C., Julicher, F. and González-Gaitán, M. (2011). Dynamics of Dpp signaling and proliferation control. Science 331, 1154–1159.

Wu, H.-H., Ivkovic, S., Murray, R. C., Jaramillo, S., Lyons, K. M., Johnson, J. E. and Calof, A. L. (2003). Autoregulation of Neurogenesis by GDF11. Neuron 37, 197–207.

Xiong, F., Ma, W., Hiscock, T., Mosaliganti, K., Tentner, A., Brakke, K., Rannou, N., Gelas, A., Souhait, L., Swinburne, I., Obholzer, N. and Megason, S. G. (2014). Interplay of cell shape and division orientation promotes robust morphogenesis of developing epithelia. Cell 159, 415–427.

Xiong, F., Tentner, A. R., Huang, P., Gelas, A., Mosaliganti, K. R., Souhait, L., Rannou, N., Swinburne, I. A., Obholzer, N. D., Cowgill, P. D., Schier, A. F. and Megason, S. G. (2013). Specified neural progenitors sort to form sharp domains after noisy Shh signaling. Cell 153, 550–561.

Zhang, H., Stallock, J. P., Ng, J. C., Reinhard, C. and Neufeld, T. P. (2000). Regulation of cellular growth by the Drosophila target of rapamycin dTOR. Genes & Development 14, 2712–2724.

